# A spatial transcriptomic atlas of acute neonatal lung injury across development and disease severity

**DOI:** 10.1101/2025.06.02.656433

**Authors:** Saahithi Mallapragada, Ruqian Lyu, Arianna L. Williams-Katek, Brandon K. Fischer, Annika Vannan, Niran Hadad, Evan D. Mee, Shawyon P. Shirazi, Christopher S. Jetter, Nicholas M. Negretti, Anne Hilgendorff, Laurie C. Eldredge, Gail H. Deutsch, Davis J. McCarthy, Jonathan A. Kropski, Jennifer M.S. Sucre, Nicholas E. Banovich

## Abstract

A molecular understanding of lung organogenesis requires delineation of the timing and regulation of the cellular transitions that ultimately form and support a surface capable of gas exchange. While the advent of single-cell transcriptomics has allowed for the discovery and identification of transcriptionally distinct cell populations present during lung development, the spatiotemporal dynamics of these transcriptional shifts remain undefined. With imaging-based spatial transcriptomics, we analyzed the gene expression patterns in 17 human infant lungs at varying stages of development and injury, creating a spatial transcriptomic atlas of ∼1.2 million cells. We applied computational clustering approaches to identify shared molecular patterns among this cohort, informing how tissue architecture and molecular spatial relationships are coordinated during development and disrupted in disease. Recognizing that all preterm birth represents an injury to the developing lung, we created a simplified classification scheme that relies upon the routinely collected objective measures of gestational age and life span. Within this framework, we have identified cell type patterns across gestational age and life span variables that would likely be overlooked when using the conventional “disease vs. control” binary comparison. Together, these data represent an open resource for the lung research community, supporting discovery-based inquiry and identification of targetable molecular mechanisms in both normal and arrested human lung development.

## INTRODUCTION

The intricate process of forming functional gas exchange units in the lung begins in utero and continues through early adolescence (1–5). Recent work has made substantial progress in dissecting the transcriptional shifts and cell populations present in both normal and disrupted lung development at single-cell resolution (6–11). From these disaggregated cellular datasets, we have learned that the molecular classification of developmental stages is more nuanced than previously appreciated from histologic assessments alone. Cell lineages mature at different rates, with transcriptional expression profiles that presage structural changes in tissue (12–15). With these technical and conceptual advances, there is a growing need to understand the spatial relationships of stage-specific specialized cell populations within the developing lung under normal and injured conditions. While single-cell RNA sequencing (scRNA-seq) datasets have provided a foundation for understanding the cellular populations during development and injury, these data lack spatial context, preventing delineation of cell localizations and cell - cell relationships and their organizational changes during organogenesis.

To address these critical questions, we profiled a cohort of human infant lung samples at varying stages of lung development, injury, and disease, using imaging-based spatial transcriptomics, generating transcriptional profiles of ∼1.2 million cells in their spatial context. Comparative analysis of human infants with variable development, chronological age/life span, and stage of disease presents a complex problem, especially when selecting the appropriate comparator samples. Preterm birth itself is an injurious stimulus to the developing lung, and injury superimposed upon development requires simultaneous consideration of both developmental stage and disease severity. Furthermore, the diagnosis and classification of chronic lung disease is vulnerable to subjective criteria and defined by the patient’s respiratory support, which is not a fixed variable. Detailed clinical metadata about longitudinal respiratory support are not always available, and the clinical staging of chronic lung disease is not aligned with molecular and structural changes in the lung (7). To overcome these challenges, we developed a new analytical approach based only on the widely obtained, objective features of gestational age at the time of birth and the chronological age/life span at the time of sample collection, with the goal to develop a strategy that could be universally applied to tissue from other lung repositories.

Here, we report the first findings from these analyses, which validate known cellular structures within the developing lung and generate new hypotheses about potential molecular drivers that may influence organogenesis with respect to life span and gestational age. Finally, to serve as a resource for the lung research and developmental biology community, we have made these data available as a public data explorer at https://lungcells.app.vumc.org/public/sucre/neonatal_spatial/. Our overarching goal is for these spatial-temporal data to serve as a foundation for hypothesis generation surrounding the cellular and molecular drivers of lung development and the identification of potential targetable molecular mechanisms, while providing an analysis strategy informed by relationships between transcript and cellular niches.

## RESULTS

### Characterizing the cellular composition of the developing human lung

To spatially resolve transcriptomic features of the neonatal lung during normal development and disease, we analyzed infant lung samples across different developmental time points using imaging-based spatial transcriptomics. This spatial dataset consists of 17 lung samples, ranging from 16 weeks gestational age (GA) to 40 weeks GA, with varying degrees of lung injury (Fig. S1, Supplemental Tab. S1). From each sample, we analyzed two adjacent 3mm x 3mm square sections arranged in a tissue microarray (TMA) using the Xenium platform (Fig. 1a, Supplemental Fig. S1). We recovered a total of 229,507,430 transcripts for analysis, including 76,379,584 partitioned into 1,170,328 cells (see Methods). From these, we identified 40 distinct cell populations (13 epithelial, 5 endothelial, 6 mesenchymal, and 16 immune), annotated using prior published hallmark genes (Fig. 1b - d, Supplemental Fig. S2, Supplemental Tab. S2).

**Figure 1.**
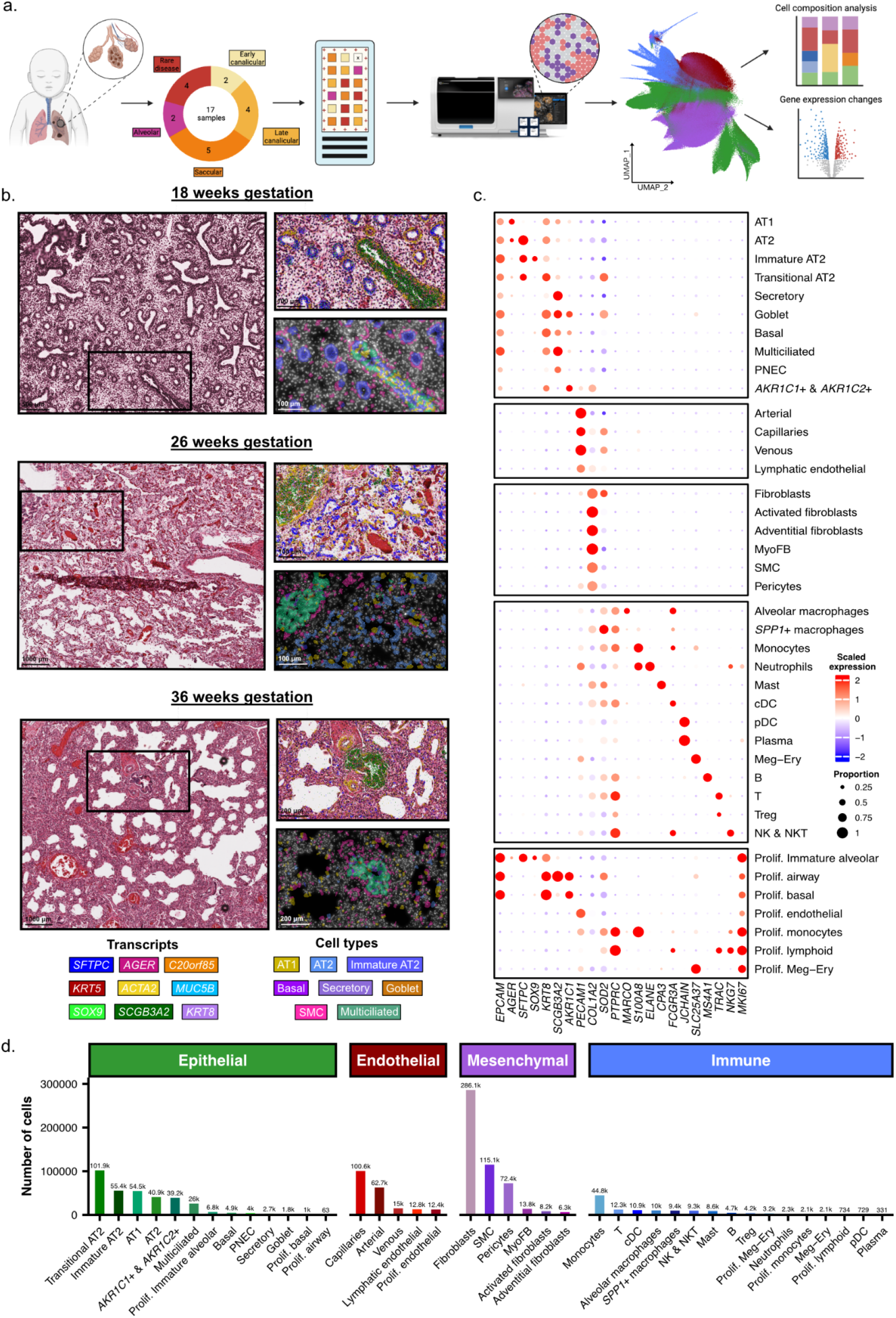
Spatial transcriptomic analysis of human infant lung tissue reveals distinct cell populations spatially localized to histological structures. a) Schematic representation of the study design. b) H&E images of infant lung tissue within this study at different developmental and gestational ages, with transcripts overlayed (top inset) colored by hallmark gene transcript, and a Xenium explorer image (bottom inset) colored by cell type. c) Dotplot heatmap demonstrating representative hallmark genes used to annotate all 40 cell types, with the diameter of each circle indicating the relative proportion of cells expressing the transcript and the intensity of color representing the abundance of transcripts. d) A bar plot demonstrating the frequency of each cell type across the entire dataset, split by epithelial, endothelial, mesenchymal, and immune cell lineages.

### Characterizing changes in tissue architecture across development

To assess cellular and molecular changes associated with development, we grouped samples into developmental groups based on gestational age at the time of birth: Early canalicular (16–18 weeks GA), late canalicular (21–24 weeks GA), saccular (26–35 weeks GA), and alveolar (36 weeks GA) (Supplemental Fig. S3). In addition, this dataset also includes samples from patients diagnosed with rare developmental lung diseases: Congenital high airway obstruction syndrome (CHAOS), pulmonary hypoplasia and dysplastic kidneys, angiotensin receptor blocker (ARB) fetopathy-induced bronchopulmonary dysplasia (BPD), and one lung explant from a 30-year-old woman who had severe BPD as an infant. As expected, we observed differences in cell type proportions across developmental groups, alongside variability in the rare disease samples (Fig. 2a, Supplemental Fig. S4). Furthermore, we identified a population of immature alveolar type 2 (AT2) cells marked by increased expression of *SOX9* alongside canonical expression of *SFTPC*, which appeared nearly exclusively in patients under 21 weeks of gestation (Fig. 2b, Supplemental Fig. S2, Supplemental Fig. S5, Supplemental Tab. S3). Enabled by spatially-aware analyses, we next moved beyond cell type classification to define the landscape of cell-cell interactions within the developmental stage-specific pattern, using cell proximity as a proxy for potential interaction. Briefly, within each developmental stage and across all pairwise combinations of cell types, we quantified the likelihood that cell type pairs were in close proximity to one another (see Methods). We focused on cell type pairs that were present in sufficient numbers among all gestational age groups and were significantly enriched/depleted for being in proximity to one another in at least one group. Importantly, this represents a comprehensive map of cell-cell interactions across the developing human lung (Fig. 2c, Supplemental Fig. S6 - S14, Supplemental Tab. S4). We identified representative interaction pairs that showed stage-specific patterns of cell proximity across the four developmental stages along with each rare diseased sample separately (Fig. 2d). As a confirmatory example, we find capillaries and alveolar type 1 (AT1) cells are highly unlikely to occur next to one another in samples from the early and late canalicular periods; however, in samples from the alveolar stage and the adult sample, they are more likely to be close to one another.

**Figure 2.**
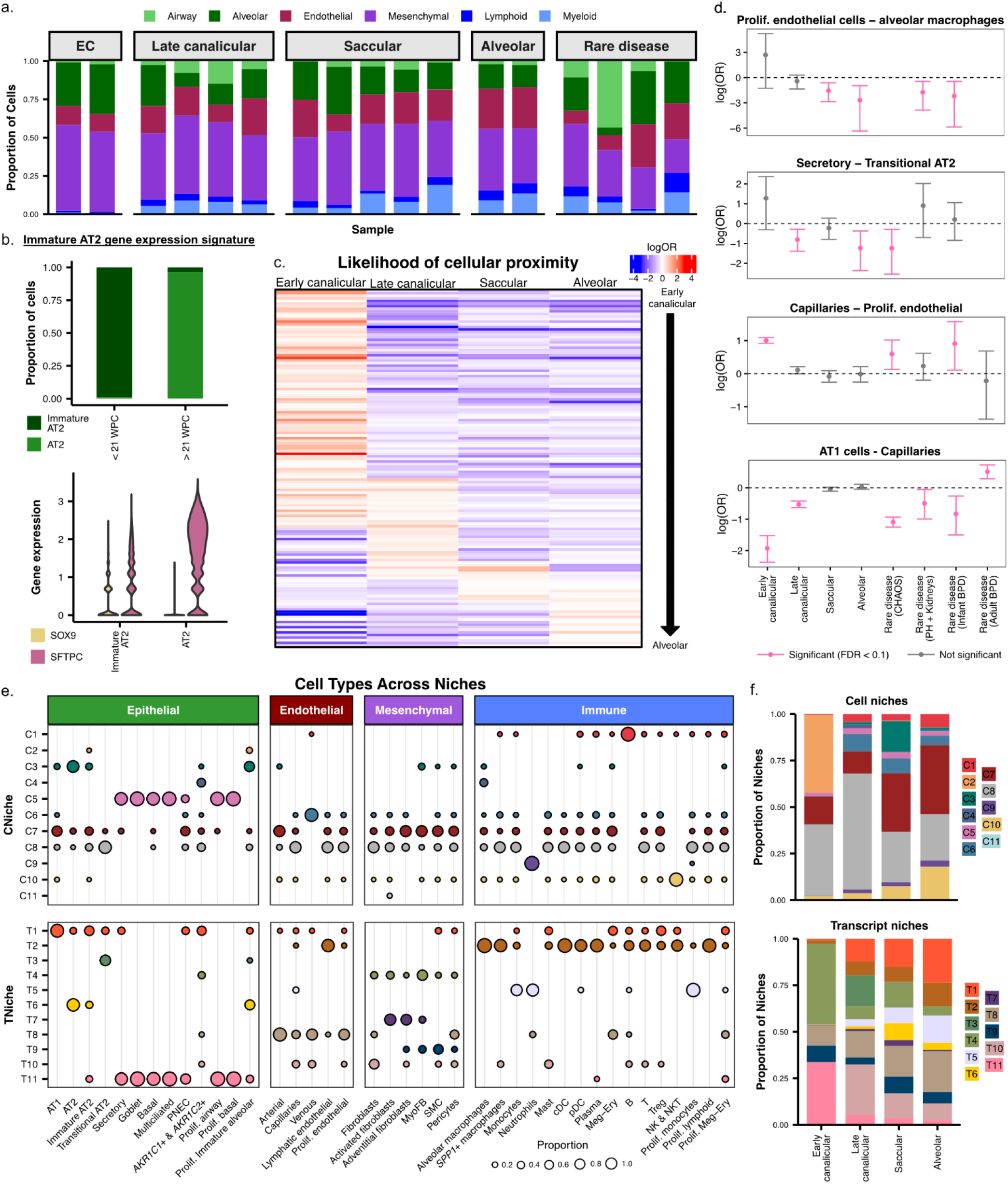
Analysis of the cellular and molecular neighborhoods and “niches” identifies a unique pattern of cell-cell interactions associated with developmental stage. a) A stacked bar plot highlighting the proportion of each sublineage across every sample in this cohort. Samples are grouped by developmental stage and rare disease. Rare diseases included (in this order): CHAOS syndrome, pulmonary hypoplasia + dysplastic kidneys, ARB fetopathy-induced BPD, and the 30-year explant with BPD. b) A barchart and violin plot demonstrating shifts in cell type proportion between AT2 and immature AT2 (top panel) and the increase of *SOX9* and decrease of *SFTPC* expression in immature AT2 cells (bottom panel). c) A heatmap of log odds ratios (logOR) quantifying proximity likelihood between cell types across developmental windows. Each row represents a cell-cell interaction probability with red indicating higher likelihood of two cells being in close proximity. Results can be found in Supplemental Tab. S4. Early canalicular to alveolar stages are ordered left to right. d) Lollipop plots showing logOR and confidence intervals for selected cell type - cell type proximity across each developmental stage and rare disease samples. In each cell type pair, the cell type listed first is considered the starting cell in the analysis; e.g., for the Secretory – Transitional AT2 pair, logOR and confidence intervals reflect the enrichment/depletion of Transitional AT2 cells as the single nearest neighbors of Secretory cells (see Methods). Pink error bars indicate statistically significant results (FDR < 0.1); gray indicates non-significance. e) The distribution of cell types across defined niches. Dot plot shows the presence and relative abundance of specific cell types within both cell-based and transcript-based niches (top: “CNiche”, bottom: “TNiche”), stratified by major lineage (epithelial, endothelial, mesenchymal, immune). Dot size reflects the proportion of cells within that cell type & niche. f) Niche proportion across developmental stage. Stacked bar plots show the composition of cell niches (top) and transcript niches (bottom) across gestation. For both CNiches and TNiches, proportions reflect the number of cells, not transcripts, assigned to a given niche.

We then examined cellular profile changes in tissue architecture more broadly, moving beyond pairwise comparisons of cell type proximity. To this end, we partitioned regions of tissue into discrete “niches”. The goal of this approach was to comprehensively characterize each tissue section and identify any cellular or molecular similarities across the cohort. Each niche represents a region of molecular or cellular similarity defined by the spatial proximity of either cells or transcripts (see Methods). Briefly, cellular niches (CNiches) were defined by creating a local neighborhood for each cell based on spatial proximity and cell type annotation, followed by k-means clustering of neighborhoods using cell type composition. Transcript-defined niches (TNiches) are conceptually similar, but utilize spatial neighborhood structures of transcripts rather than cells and cell type annotations. Once TNiches are defined, cells can be assigned TNiche labels based on the unified spatial coordinates. (see Methods). Across both approaches, we identified 11 multicellular niches (Fig. 2e, Supplemental Fig. S15). Both approaches find generally concordant results, but inherently partition the tissue slightly differently. A representative example would be TNiche 1, which represents the alveolar niche and is composed of the key alveolar cell types, including AT1, AT2, and capillaries (Fig. 2e). As further confirmation that these computationally-defined niches represent biologically relevant regions within the lung, we observe that TNiche 1, in particular, increases in abundance across gestation, consistent with the cell type proximity analysis and alveologenesis throughout lung development (Fig. 2f, Supplemental Fig. S16). Additional niche variation across gestational age was also observed (Fig. 2f).

### Gene expression changes associated with development and neonatal injury

Characterization of the cellular landscape was complemented by the identification of gene expression changes associated with both development and neonatal lung injury by pathologic staging. Prior work characterizing dysregulation of gene expression in neonatal lung injury and BPD has classified individuals into discrete groups such as evolving BPD (neonatal lung injury prior to 36 weeks gestational age), BPD, and controls (16,17). However, these groupings, particularly evolving BPD, are often difficult to adjudicate and do not capture the full spectrum of heterogeneous injury. Building upon the idea that every infant born prematurely experiences some level of neonatal lung injury, we postulated that life span may serve as a proxy for the degree of neonatal lung injury (Fig. 3a). However, given the rapid developmental changes occurring from the canalicular to alveolar stages, we also anticipated life span would need to be considered within the context of gestational age. To this end, we quantified the effects of both life span and gestational age on gene expression. Briefly, for 11 samples ranging from 21 - 40 weeks GA at the time of birth and 0 - 16 weeks of life span (Fig. 3b), we tested for changes in gene expression within each cell type using a linear regression (see Methods). Our regression model included terms for both gestational age and life span, allowing us to identify gene expression changes associated with life span accounting for the effects of gestational age and vice versa. We focused our expression based analyses on structural cells as we found the immune compartment to be particularly subject to spurious associations driven by transcript contamination from mesenchymal cells (see Discussion; Fig. S17; Tab. S6 - 7). In total, we found 25 gene expression changes associated with life span and 18 associated with gestational age (Fig. 3c - d, Supplemental Fig. S18, Supplemental Tab. S8; FDR < 0.1; see Methods).

**Figure 3.**
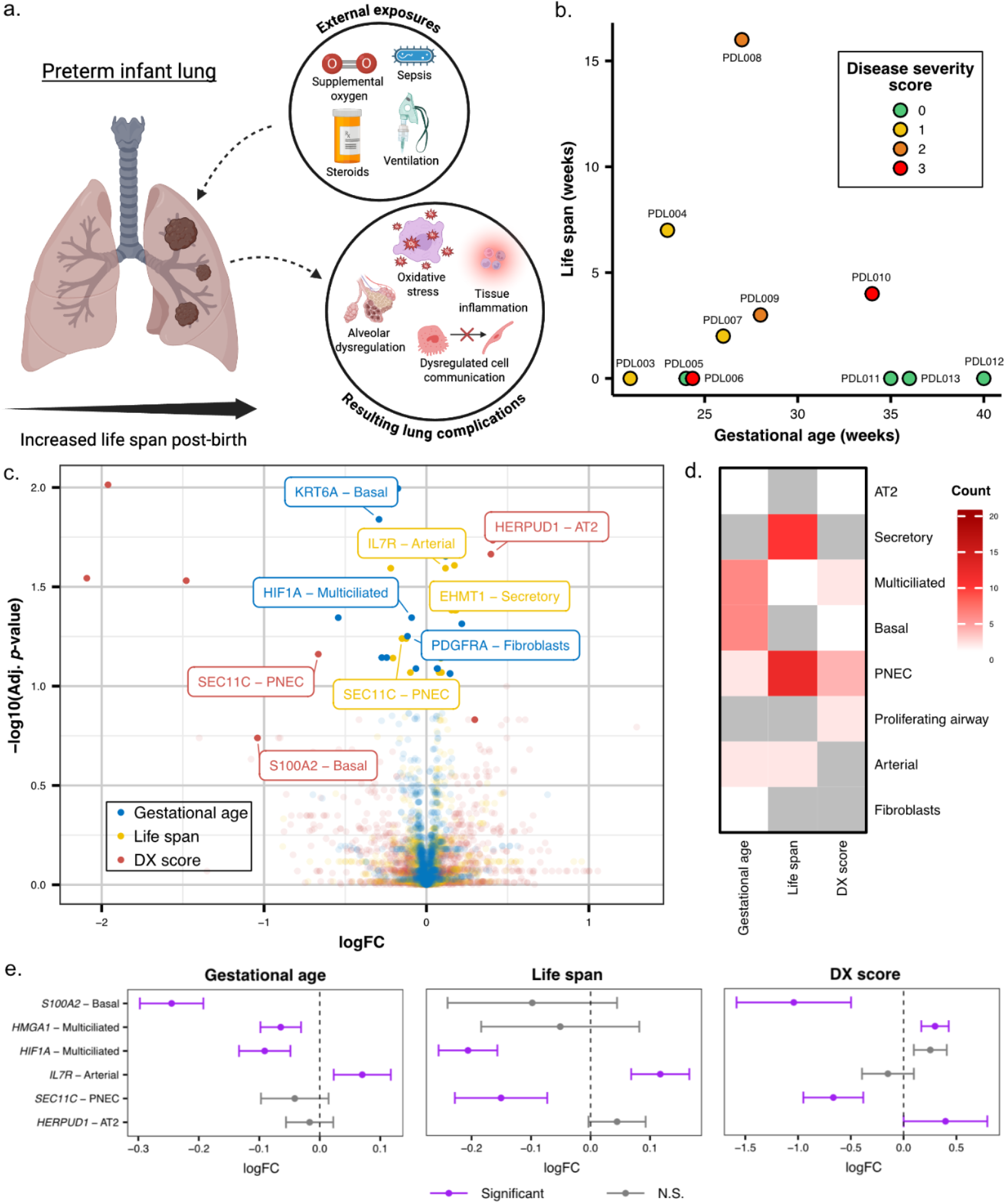
The impact of gestational age, life span, and disease severity on lung biology in preterm infants. a) An overview of how gestational age and environmental factors can alter lung development. The graphic illustrates how supplemental oxygen, ventilation, sepsis, and other external factors (often used when an infant is born early) can contribute to alveolar dysregulation, oxidative stress, and tissue inflammation, ultimately disrupting cell-cell communication and overall lung development in preterm infants. b) A scatterplot showing each sample’s gestational age (x-axis) and postnatal life span (y-axis), with points colored by disease severity (DX) score (0–3). c) Volcano plot of gene expression associated with gestational age, life span, and disease severity score. Log fold change (logFC) plotted against FDR for genes significantly associated with at least one variable. Labeled genes are color-coded by the associated condition: gestational age (blue), life span (yellow), and disease severity score (red). d) Gene - cell type associations between gestational age, life span, and disease severity. The heatmap displays the number of significantly associated genes per cell type for each variable. Higher counts (darker red) indicate a higher number of significantly differentially expressed genes in that cell type. e) Significant genes that are upregulated in specific cell types between gestational age, life span, and disease severity score. Lollipop plots show logFC and confidence intervals for select gene - cell type pairs across gestational age, life span, and disease severity. Purple indicates significant associations (two-level FDR cutoff, see Methods); gray indicates non-significant (N.S.).

As an orthogonal approach, we worked with a pediatric pulmonary pathologist to create a disease severity (DX) score (see Methods) for every tissue section in our study. This score represents the overall degree of damage observed in the tissue, aggregated across multiple features. When comparing the molecular data to histologic data (length of life span and gestational age) of each infant, the DX score assigned accurately matches the molecular data being measured. Using a similar regression framework (see Methods) and again focusing on structural cells (Tab. S7), we identified 10 genes associated with disease severity (Fig. 3d, Supplemental Fig. S18, Supplemental Tab. S9). Intersecting results from associations with gestational age, life span, and disease severity (see Methods), we found 7 gene-cell type pairs overlapped between gestational age and life span; 12 are shared between gestation age and DX score; 2 between life span and DX score; and 1 gene-cell type pair overlaps all three terms (Fig. 3b, 3c, Supplemental Fig. S18). Select gene-cell type pairs that overlap between gestational age, life span, and DX score were chosen to showcase the expression differences (in log fold change, logFC) between comparators (Fig. 3e).

## DISCUSSION

This atlas provides a resource to begin decoding the molecular and cellular dynamics and spatial relationships of changes in human lung development, injury, and repair. With this dataset, we gain insight into the effects of premature birth and injurious responses early in life. Our spatial transcriptomics analysis preserves the anatomical and microenvironmental context of these cell populations, providing new insights about tissue organization and the link between spatiotemporal transcriptional changes and structural injury previously prohibited by disaggregated single-cell transcriptomic analysis alone.

As with any new technology, the first step of validation with known biological controls builds confidence in future findings. Across all organs and platforms, cell segmentation remains a challenge in spatial transcriptomic analysis. This is especially true in the lung where proximally, cells in the pseudostratified airway epithelial appear to “overlap” each other by 2D section, and distally where AT1 cells and aCaps have extensive, flattened cellular dimensions directly adjacent to each other (18). Despite this caveat, we were reassured that our methods identified 40 distinct cell populations with hallmark genes consistent with prior published work in humans and other mammalian systems (6,8,15,19,20). Indeed, across lung development, we identified cell types with transcriptional profiles that are not only stage-specific, but also with transcriptional profiles associated with exposure to environmental stimuli (i.e., supplemental oxygen and mechanical ventilation). A particular strength of this approach is the number of cells recovered from the samples and the ability to match the molecular and cellular data with histologic features. The number of cells recovered is much greater than all previously published human neonatal scRNA-seq datasets combined (∼100,000 cells per sample via spatial vs. ∼5,000 cells per sample via scRNA-seq). This abundance of cells allows us to identify and annotate rare cell types and study their associations with neighboring cells and niches. Additionally, we present data here that can aid in our collective understanding of transcriptional differences in known cell types - differences that can now be linked to developmental stage and chronological age.

Historically, the stages of lung development have been delineated based largely on the pattern of airspace formation observed on two-dimensional histologic sections (3,4). Here, we find that there is a reproducible pattern of cell-proximity relationships linked to developmental stage. How the epithelial cells associate with endothelial cells to form a functional alveolus is an area of active investigation, with recent live-imaging data overturning the notion of ingrowing septation in favor of a model of outward-growing alveolar epithelial cells supported by a mesenchymal ring structure (21). Within our spatial dataset, we find data supportive of AT1 cells and capillaries gradually becoming closer in proximity as lung development progresses. In addition, these data generate multiple hypotheses for future testing.

The comparison of human tissue samples relies heavily on accurate grouping by disease status. In diseases with clear biomarkers or pathologic hallmarks of diagnosis, e.g., cancer, pneumonia, this is a relatively straightforward endeavor. In the case of lung injury superimposed on lung development, which comparator tissue to use remains a scientific challenge. Foremost, all preterm birth represents an injury to the developing lung, as even exposure to room air and physiologic distension pressures are in excess of the oxygen levels and pressures experienced in utero (22). Further, neonatal biopsies and autopsies are rare, as are lung transplants, making this group significantly underrepresented in most large aggregated human cell atlases. This is compounded by the diagnosis of BPD, the leading cause of chronic lung disease in infants, which is defined by respiratory support received and most certainly comprises multiple clinical/anatomic endotypes (23,24). With a goal of circumventing the inherent pitfalls of imposing binary “disease vs. control” comparison on a disorder with a spectrum of manifestations, we instead created an analysis framework reliant only upon objective data that could be applied to every sample, e.g., gestational age and life span. We therefore propose a paradigm shift in the classification of neonatal tissue by objective categories that moves away from binary disease/control categories.

Beyond eliminating the bias and challenge of grouping samples into disease and controls, this approach has several strengths, including the ability to be applied to smaller sample sizes typically found in pediatric repositories or in rare disease settings. From this framework, we have identified that gestational age and life span at the time the sample is obtained can independently shape gene expression and perturb developmental pathways. Moreover, our data support the notion that while lung maturation can be a function of both gestational age and life span, the transcriptional profiles of ex-utero development do not necessarily align with in utero development and that in fact, prolonged exposure of a preterm lung to oxygen and mechanical ventilation may correlate with transcription of some genes associated with gestational age, e.g., expression of *HIF1A* by multiciliated cells or *IL7R* by arterial endothelial cells (Fig. 3E).

There has been a tendency to classify the lung injury and abnormal development associated with preterm birth as representative of senescence or developmental regression, with each conclusion supported by the expression of representative hallmark genes associated with each process (25,26). While we understand the need to classify newly discovered profiles within a heuristic of previously described processes, the grouping of dissimilar processes based on the presence of a few transcripts may create more confusion than clarity. Future work with an expanded library of samples using this more agnostic approach may provide more insights into how the abnormal development associated with preterm birth is both similar and different to normal development and senescence. Notably, the identification of shared gene-cell type pairs across developmental stage, chronological age, and disease score may highlight molecular drivers that have a shared or opposite effect across these three clinical variables, with a goal of identifying targetable regulators to mitigate injury-induced dysregulation or promote normal development.

Some limitations of this study hindered our ability to conduct deeper analyses. First, while our dataset is large in terms of cells analyzed, the number of distinct donors remains limited, and this limitation is more striking when the donors are stratified by developmental stage, disease type, or injury severity, limiting the statistical power for detecting cell-type specific or gene expression changes by condition. In addition, technical limitations of cell segmentation inherent in spatial transcriptomics can result in contaminating transcripts from adjacent cells being misclassified as belonging to adjacent cells. Indeed, immune cells being assigned transcripts representing hallmark genes from parenchymal cell types represented a technical challenge throughout the dataset. We suggest that unlike other parenchymal cells, immune cells are generally surrounded by non-immune cells in the tissue making them more likely to have contaminating RNA present within the field of their nucleus.This can be particularly impactful when performing differential expression analyses anchoring on biological variables which are also associated with tissue remodeling (e.g., gestational age). Even with stringent filtering approaches (See Methods) we found nearly all differentially expressed genes identified in immune cells appeared to be driven by contamination from mesenchymal cells – leading us to remove immune cells from our differential expression analysis (Tab. S6 - S7).

As such, we remain circumspect in drawing large conclusions from these data, but instead provide a robust framework for analysis applicable to small cohorts, with a goal of expanding this work to larger sample sizes from the neonatal population. The panel of genes reported here was created to apply generally to the lung, but not specific to organogenesis or neonatal lung injury. We envision that ongoing and recently published work identifying genes of particular relevance to development, injury, and repair will inform the curation of future gene panels, improving the ability to discern expression differences across conditions. Finally, while gestational age and life span are objective, the assignment of disease severity score relies upon pathological classification that may introduce an element of subjectivity.

Taken together, the data from these analyses provide a spatially-resolved map of human neonatal lung development and injury at single-cell resolution, and make these rare specimens widely available to the research community. Here, we begin to untangle how cellular composition, spatial organization, and molecular signaling are perturbed by premature birth and evolving disease pathology, and offer a benchmark for future spatial analyses aimed at identifying targetable therapeutics for preterm infants that will restore normal transcriptional profiles and structural lung development.

## Data availability

Raw and processed data are deposited at GEO under accession GSE297945. Spatial cell atlas is available at https://lungcells.app.vumc.org/public/sucre/neonatal_spatial/.

## Code availability

Custom R and Python scripts for this project are available on GitHub at https://github.com/Banovich-Lab/A-spatial-transcriptomic-atlas-of-acute-neonatal-lung-injury.

## Acknowledgements

This work was supported by National Institutes of Health (NIH) R01HL145372 (to J.A.K. and N.E.B.), U01HL175444 (to J.A.K., N.E.B. and J.M.S.S.), R01HG011886 (to N.E.B. and D.J.M.), K08143051 (to J.M.S.S.), R01HL168556 (to J.M.S.S.), National Health and Medical Research Council (NHMRC) (GNT1195595 and GNT1162829 to D.J.M.), the transregional collaborative research center PILOT - TRR 359 Perinatal Development of Immune Cell Topology, Project number 491676693, funded through the Deutsche Forschungsgemeinschaft (DFG, German Research Foundation) (to A.H).

## METHODS

### Subjects and sample nomenclature

Formalin-fixed paraffin embedded lung tissue from 17 subjects from within the Human Infant Lung Repository, a resource of lung tissue from across lung development comprising pediatric autopsy and biopsy specimens (27). The use of Human Infant Lung Repository tissue has been reviewed and approved by the Vanderbilt University Institutional Review Board. Clinical metadata can be found in Supplemental Tab. S1. Sample names were generated by appending ‘PDL’ and then ranged from ‘001’ - ‘017’ (ex. PDL001).

### Tissue microarray construction

A tissue microarray (TMA) was built in order to optimize tissue capture on an entire Xenium slide (10.45mm x 22.45mm) and analyze multiple samples at once. A 5μm section of each lung sample formalin-fixed paraffin-embedded (FFPE) block was H&E stained and presented to a clinician, who then picked specific areas in each sample that were of interest. One core from each sample was manually punched out using the “Quick-Ray Manual Tissue Microarrayer Full Set” and a custom square-shaped 3mm tip. The square tip was designed and manufactured at Arizona State University’s Instrument Design and Fabrication Machine Shop (28). To that note, a custom square mold was created, and paraffin wax was poured to mold a custom recipient block with empty core spaces for sample core placement. After this, the block was placed on a clean slide and melted slightly (>2min at 60°C) to even the top out and melt the cores into the paraffin block. Finally, the block was cooled at room temperature overnight, sealed in a dry plastic bag, and stored at 4°C.

During the construction of this TMA, many of our samples were too close to one another and ended up touching. This was because the custom mold we had made created a wax grid that had extremely thin walls, so some sample cores were not as rigid and shifted position slightly as the block was melted together.

### Xenium processing and workflow

Currently, all imaging-based spatial transcriptomics technologies require the use of designated gene panels. We built the gene panel for this dataset based on an early version of the Xenium human lung base panel (PD_336) and a custom panel that we designed based on canonical cell type markers, known developmental progenitors, and previously-established lung datasets (19). 246 genes came from their base panel, while 97 genes came from the custom panel, resulting in 343 unique genes. A number of probes were assigned to each gene that contained two paired sequences that were complementary to the targeted RNA molecule, alongside a gene-specific barcode. Those paired ends are ligated, which allows amplification of the now-circular probe and bolsters the signal-to-noise ratio needed for optimal target detection and decoding.

Before any sample preparation began, all workstations were cleaned using RNase Away (RPI 147002) to remove any potential remnant RNase molecules, followed by 70% Isopropanol to remove the residue and any potential contaminants. All reagents used in this protocol were molecular grade nuclease-free. To prepare the samples for Xenium processing, we sectioned the constructed TMA on a microtome (Leica RM2135) into 5μm sections and rehydrated each slice. Two adjacent sections from the block were taken, creating two technical replicates per sample. The two chosen slices were placed on separate Xenium slides, left to dry overnight, and then stored in a sealed desiccator at room temperature (can be stored for up to 10 days). After this, each slide was placed into an imaging cassette, and we then conducted tissue deparaffinization and decrosslinking steps to make all RNA molecules targets and accessible. Each slide was then hybridized with the probes assigned to the gene panel we had previously built, which happened overnight for 18 hours at 50°C (Bio-Rad DNA Engine Tetrad 2). Following that, we removed unbound probes from the slides through subsequent rounds of washing. We then added ligase to circularize the paired ends of bound probes, which we did for 2 hours at 37°C, and then we conducted enzymatic rolling circle amplification for another 2 hours at 30°C. We then washed with PBS and PBS-T, chemically quenched autofluorescence, and used DAPI to stain all nuclei present.

Once all protocols had finished, we stored both slides in PBS-T at 4°C (can be stored up to 5 days) in a dark environment until they went on the Xenium Analyzer instrument. All Xenium FFPE preparation guidelines and buffer recipes can be found in 10X Genomics’ Demonstrated Protocols CG000578, CG000580, and CG000749.

The Xenium Analyzer instrument is a complex machine that images tissue samples in multiple cycles to accurately decode subcellular localized RNA target molecules and assign them to specific genes. In order to do this, we first marked regions of interest on the slide by manually selecting where each sample was located on each slide. One Xenium run can process up to two slides at a time, so both sections we had processed were imaged at the same time as demonstrated in Supplemental Fig. S1. We then loaded in consumable reagents, and internal fluid mechanisms controlled how those reagents were utilized and distributed. Through the course of the experiment, the machine conducts a repeated cycle of fluorescently-tagged probe binding, image acquisition, and subsequent probe stripping. These fluorescently-tagged probes were imaged at a resolution of 4240 × 2960 pixel fields of view (FOV). This cycle repeats, and after the entire process, each localized point of fluorescent intensity that was detected (RNA puncta) presented a unique pattern. Each gene on our panel was assigned to one of these unique patterns, so puncta that matched one of those patterns were decoded and assigned to that corresponding gene label. The last step in this process involved computationally stitching together all FOVs and assigned transcripts to the DAPI-stained image, in order to spatially localize where each RNA transcript is in relation to nuclei. The data in this paper were generated on software version 1.6.1.0 and analysis version xenium-1.6.0.8. Documentation regarding the manufacturing process and instrumentation can be found in 10X Genomics’ Demonstrated Protocol CG000584.

### Post-run histology and staining

Finally, immediately after both slides had finished running, we pulled them off of the Xenium Analyzer instrument and performed H&E staining to provide a comprehensive anatomical visualization of each sample. To do this, first slides had the autofluorescence quencher removed using Sodium Hydrosulfite according to 10X Genomics’ Demonstrated Protocol CG000613. We stained each slide with the following protocol in order: circulating DI water for 1 minute, hematoxylin (Biocare Medical CATHE) for 1 minute 10 seconds, DI water for 3 minutes, bluing solution (Biocare Medical HTBLU-M) for 1 minute, DI water for 1 minutes, 95% alcohol for 1 minute, eosin (Biocare Medical HTE-GL) for 20 seconds, 95% alcohol for 1 minute, 100% alcohol for 30 seconds (2 washes), and xylene for 3 minutes (2 washes). Finally, we performed coverslipping using Micromount (Leica 3801731), which we then stored at room temperature in a dark environment to allow it to harden. We took images of each sample post-staining on a 20X Leica Biosystems Aperio CS2.

### Nuclear segmentation

Nuclei were segmented based on DAPI staining using the 10X Genomics Xenium onboard software. As in our previous work (29), we chose to use the nuclear boundaries to define cells in lieu of an arbitrary cellular expansion. Transcripts were assigned to a “cell” if they overlapped the nuclear boundary. Importantly, this boundary captures transcripts in 3D space (a cylinder rather than a sphere) meaning transcripts in both the cytoplasm and nucleus are encapsulated.

### Pre-processing workflow

#### Quality control - Upstream filtering

Xenium output resulted in both transcript information and metadata per sample. The transcript files contained a variety of metrics per each transcript, including x - y coordinates, the cell to which they were assigned (cell_id), quality value (QV), and nuclear overlap (overlaps_nucleus) - low quality transcripts (QV < 20), transcripts with overlapping nuclei (overlaps_nucleus = 1), and transcripts with a cell_id of “UNASSIGNED” were removed from further analysis. We then used the Seurat package to conduct further quality filtering on the nuclear gene expression data (30). We created gene matrices per sample based on the remaining transcripts that were partitioned into a segmented nucleus, and with those matrices and metadata information, generated Seurat objects per each sample that were merged into a singular object. We then kept nuclei that had the following metrics: >10 transcripts total, >5 unique transcripts, <75μm for nuclear area, <15% of negative codewords were detected, and 0% of negative probes were found. The percent of negative codewords per cell id was calculated by dividing the aggregated count of transcripts with the assignment “Negative codeword” with the total transcript count, while the percent of negative probes per nucleus was calculated by dividing the aggregated count of transcripts with the assignment “Negative probe” with the total transcript count, respectively. From here on out, these nuclei will be referred to as cells.

The TMA construction of these samples resulted in certain samples pressing into one another for both slides. In light of this, Xenium generated multiple files containing information from multiple samples that needed to be processed, aggregated together, and then split according to sample in order to ensure that the correct information was being assigned to the correct sample. All these data were processed using Seurat V5 (31) in R. The Xenium Explorer includes a feature that allows for freehand drawing and manual selection of specific cell ids, so we visually selected cells that we felt belonged to each sample, being careful to not include any that were located too close to another sample. This allowed us to properly separate the samples that had merged together during pre-processing and partition cells into their appropriate sample. We then removed all cells from our Seurat object that were not selected within these freehand drawings.

#### Coordinate adjustment and dimensionality reduction

All data objects were transformed into Seurat objects, and merged together into one Seurat object. All of the cells analyzed in this study were compiled from two different slides, which allowed for shared coordinates among transcripts. This resulted in overlapping samples between both slides, so each coordinate was manually adjusted to ensure proper visualization when plotting. These adjusted coordinates were then added to the Seurat object as a Dimension Reduction object.

#### Clustering, cell type annotation, and downstream filtering

We required the use of a graphical processing unit to do any sort of clustering, given the magnitude of data (32). We used a container with the rapids (v21.8.1) (33,34) - single cell Python library (ScanPy - v1.8.1) (35) to perform clustering and dimensionality reduction, and Seurat for remaining downstream analyses. We used log1p transformation to normalize gene expression, calculated principal components that were used in the dimensionality reduction, and used the Leiden algorithm to cluster cells together by computing cellular neighborhoods (36). Uniform Manifold Approximation and Projection (UMAP) plots were used for visualization (37).

For cell type annotation, we utilized the Seurat V5 FindMarkers (38) function to identify the highest expressing genes per cluster. We then used a combination of canonical genes per cell type (12,19,29) and known developmental progenitor genes (1,7,12) to anchor ourselves and segregate each cluster into 6 sublineages: airway epithelial (*EPCAM, SCGB3A2, KRT5, MUC5B, AGR3)*, alveolar epithelial *(EPCAM, SFTPC, AGER, KRT8)*, endothelial (*PECAM1, EPAS1*), mesenchymal *(COL1A2, DCN, LUM, ACTA2)*, lymphoid *(PTPRC, BANK1, CD1C, TRAC, IL7R)*, and myeloid *(PTPRC, FCER1G, JCHAIN, S100A8, MARCO)*. New Seurat objects were created based on these groupings, and processed through the same clustering workflow as described above. Within each of these objects, initial cell type labels were assigned based on FindMarkers results and more canonical genes. Specific hallmark genes used to annotate each cell type are located in Supplemental Tab. S2. We observed clusters with overall low gene expression patterns, and these clusters typically corresponded to low RNA counts. We then conducted downstream filtering and removed them from our analysis as a whole. Once all lineage-level objects were labeled, they were merged back together. To specify and separate certain cell types together, certain cell types were isolated into a new Seurat object and put through the workflow once again (29). The final level of cell type annotation resulted in 40 cell types total across the 4 lineages.

### Downstream analyses

#### Differential gene expression

We used the Limma (39) package in R to identify differentially expressed genes between the Immature AT2 and AT2 cell types across all samples - all differentially expressed genes and coordinating statistics available in Supplemental Tab. S3. In order to treat both technical replicates per sample as one unique individual, we pseudobulked across individual. We did this by aggregating all cells from both technical replicates together so that each individual has a single gene expression value per cell type. We then adjusted the *p*-value with an FDR correction and established a significance threshold of FDR < 0.1.

#### Cell-based niches

Seurat v5’s BuildNicheAssay (40) function was used to group cells into 11 spatial niches using k-means clustering based on cellular neighborhoods consisting of 10 nearest neighbor cells.

#### Transcript-based niches

We applied a Graph Neural Network (GNN) pipeline to directly analyze the spatially identified transcripts without nuclei or cell boundaries, the same approach as in a previous study (29). The GNN model was applied to learn the spatial neighbourhood structures of individual transcripts through sampling and aggregating information across neighbours. We built transcript-based graphs for individual tissues, after filtering out low-quality transcripts (qv < 20) and controlled transcripts with transcripts as nodes. Edges were added between nodes if their spatial distances were within 3 μm. We removed small connected components (size <10) in graphs, which were likely due to background signals. To jointly analyze the 34 tissues, we sampled subgraphs and merged them into a joint graph for model training. To generate the subgraph, for each tissue, we began by randomly selecting 5,000 root nodes. Then we collected their 3-hop neighbours, i.e., neighbours reached by walk length 3. Finally, we included all existing edges among the selected nodes to complete the subgraph. The subgraphs were then joined (total nodes 11,418,874, total edges 254,286,407) and used as input for training a GNN model. Specifically, we constructed a 2-layer GraphSAGE model (41,42), with both hidden layer sizes as 50, and we adopted the attentional aggregation method (43) for pooling information across spatial neighbours as implemented in StellarGraph (v1.2.1). For each focal node, we sampled 20 and 10 neighbours from its first hop and second hop of neighbours, respectively, for aggregation. The model was trained in an unsupervised way by solving a binary edge prediction task. In other words, the model parameters were tuned by learning to predict whether two nodes were connected in the input graph through similarity of their spatial neighbourhood structure after pooling (41). We then applied the trained model to all input transcripts from each tissue and obtained the latent embedding representations of individual transcripts. We applied a Gaussian Mixture Model classification on the embeddings to derive the 11 TNiches.

#### Transcript-based niche plots and assignment of cells to niches

The TNiche plots for individual tissues were created by summarizing transcript labels into hex bins with a width of 5, and hex bins with fewer than 10 transcripts were excluded. Each hex bin was labelled by the majority niche label among all the transcripts included in the same bin. TNiche labels were transferred to cell centroids. Nuclei centroids were assigned to hex bins by spatial proximity, and each centroid was assigned to its closest hex bin, therefore transferring the niche labels of the hex bins to the nuclei centroids.

#### Cell-type proximity analysis

To identify spatial relationships between cell types, we designated each cell as an anchor and calculated the distance and angular orientation to neighboring cells within a fixed 60-µm radius. We considered only immediate neighbors within this radius as proximal. To assess the likelihood of specific cell types being spatially associated, we used Fisher’s exact tests to calculate odds ratios, allowing us to categorize neighboring cells as either proximal or non-proximal and using cell type as a covariate. We first performed this analysis across all samples and cell types. Then, we grouped 13 samples in our cohort into the following developmental stages: early canalicular (16 - 18 weeks gestation), late canalicular (21 - 24 weeks gestation), saccular (25 - 35 weeks gestation), and alveolar (36 weeks gestation - term), and we partitioned the remaining 4 samples into their own “rare disease” group. We performed this analysis again within each of these groupings to identify cell type - cell type interaction pairs significantly associated with either gestational age or disease. Log odds ratios (logORs) for each group are found in Supplemental Tab. S4.

We pulled the logORs previously calculated for early canalicular (EC), late canalicular (LC), saccular (S), and alveolar (A), and binarized the data. We then used the Cutree (44) package in R to calculate a distance matrix and cluster the data, which revealed 16 unique patterns across the 4 stages of development (ex: EC → +, LC → -, S → -, A → -). We arranged these patterns in a descending stepwise manner, which identified cell type interaction pairs that were significantly associated (FDR < 0.1) with each developmental time point. We downsampled to interaction pairs that were present in all 4 developmental stages and significant in at least one.

#### Gestational age and life span regression analysis

We built a linear model with terms for gestational age and life span to determine specific genes in each cell type that have a significant association with either gestational age, life span, or both. We excluded 6 samples from our cohort - two samples that were under 21 weeks gestation and four samples that had rare diseases. We excluded these samples for this analysis because our primary objective was to assess the effect of gestational age and life span on evolving neonatal lung injury. We pseudobulked across individuals as we did for earlier analyses. We then created a filtering threshold and kept genes if they had expression in > 20% of cells in that particular cell type (29,45).

#### Pathology grading and disease regression analysis

We sought to employ the same kind of regression model, but across disease severity. Pathologic review of 11 of the 17 lung samples used for the TMA was performed by an experienced pediatric pulmonary pathologist at Seattle Children’s Hospital. Assessment was performed on the whole slide scanned at 20x as well as region of interest (ROI) used for the TMA. Clinical data known at time of review included gestational age at birth and subject age at demise. A cumulative score of disease took into context the degree of acute/subacute injury (hyaline membranes, inflammation, hemorrhage), alteration of alveolar architecture (hyperinflation, simplification) and degree of chronic remodeling (increased lobular smooth muscle, fibrosis). Vascular changes (arterial wall hypertrophy, thrombi) were also noted. The scoring system of disease was as follows: 0, no findings; 1, mild findings; 2, moderate findings; 3, severe findings. We then ran a linear model with one term for disease severity and identified genes in distinct cell types that had a significant association with pathology.

#### Intersecting between gestational age, life span, and disease severity score

We sought to find the significant gene associations that overlap across all three terms. To account for incomplete power we used a two-level FDR cutoff. Associations with a 10% FDR in each term independently were recorded, and if those associations were present in another term with a 20% FDR, those were considered to overlap between terms. For example, if a gene - cell type pair was significant (FDR < 0.1) with respect to gestational age, and has an FDR < 0.2 with respect to life span, that pair would be considered overlapping.

