## Supplemental figures S1 - 18 for "A spatial transcriptomic atlas of acute neonatal lung injury across development and disease severity"

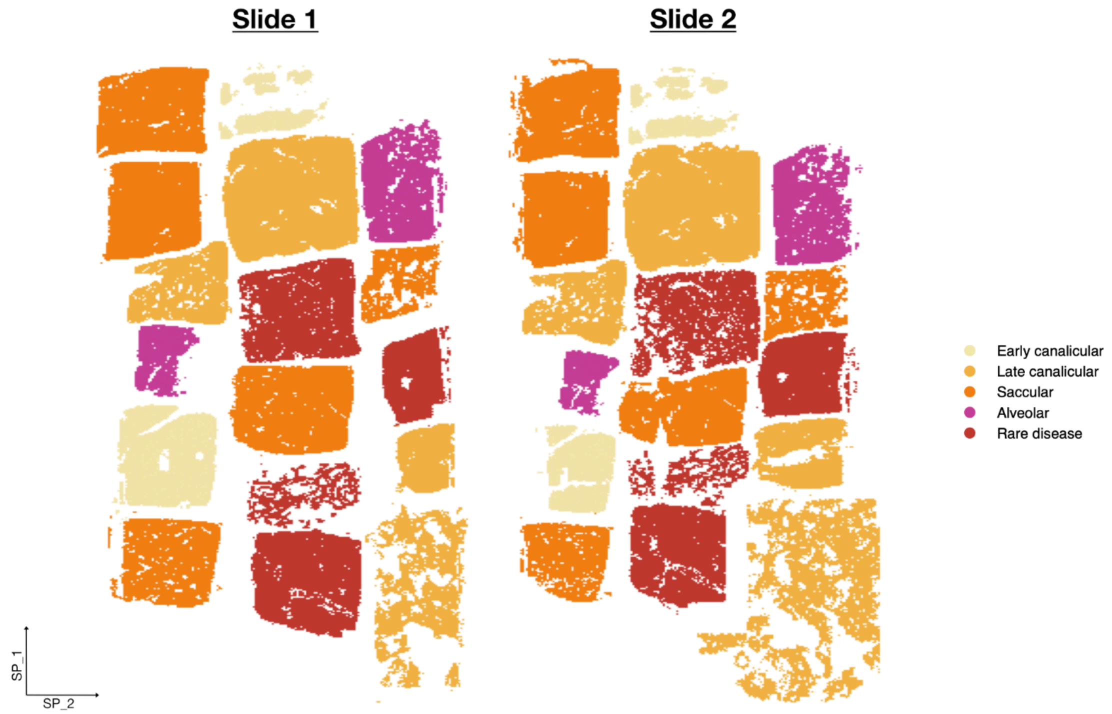


Supplemental Fig. S1:

A plot that shows each cell within each sample, highlighting the tissue structure of each sample across both slides. Each sample is colored by their gestational age and/or disease pathology.

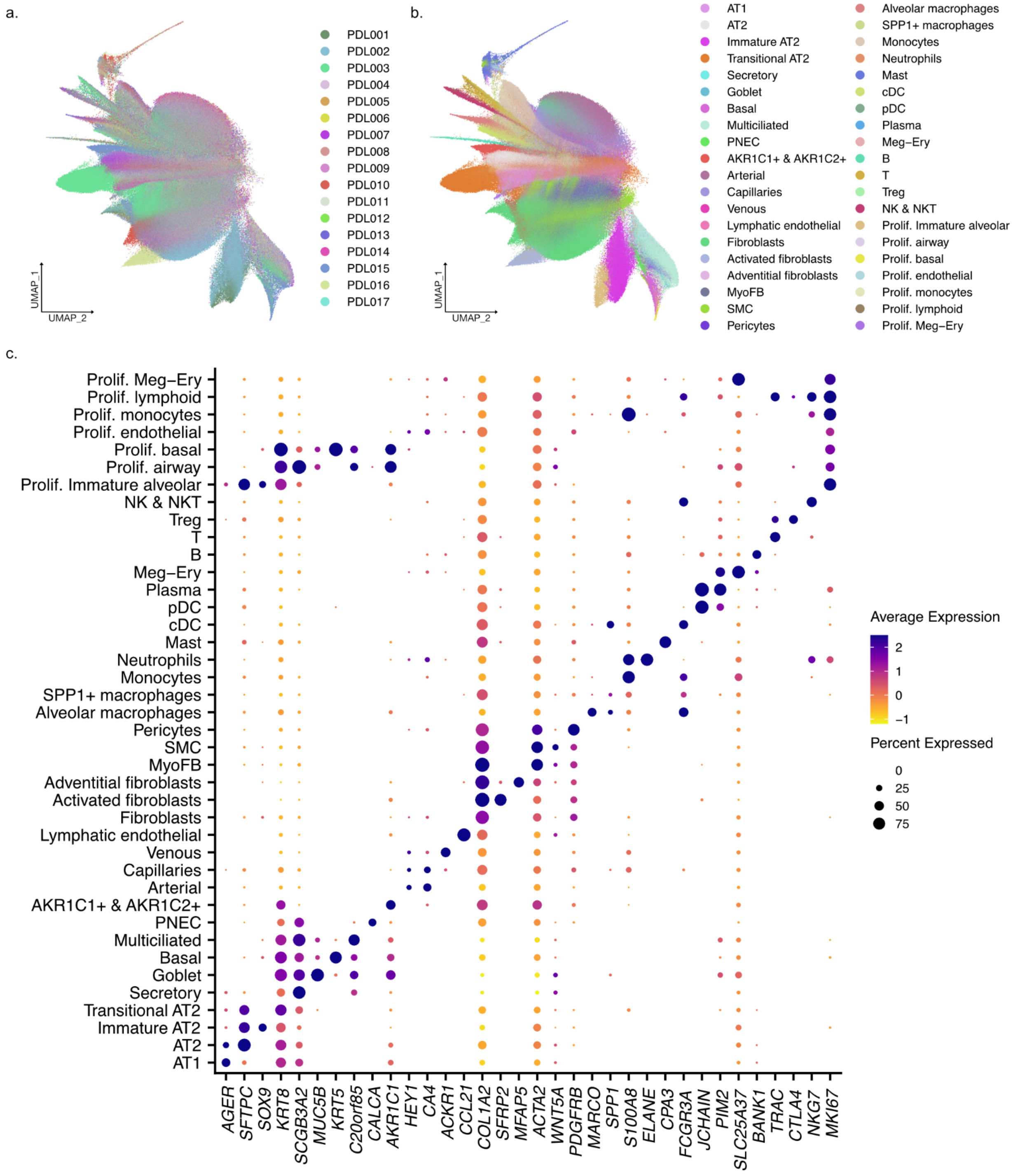


Supplemental Fig. S2:

a-b) UMAP plots showing the distribution of cells from each individual (a) and of each cell type (b). c) A dotplot highlighting hallmark genes used to annotate each cell type. Note - one cell type (*AKR1C1+ & AKR1C2+)* was labeled as such because of its high expression of AKR1C1, AKR1C2, and general airway epithelium markers (*EPCAM*, *SCGB3A2*, *SCGB1A1*, etc).

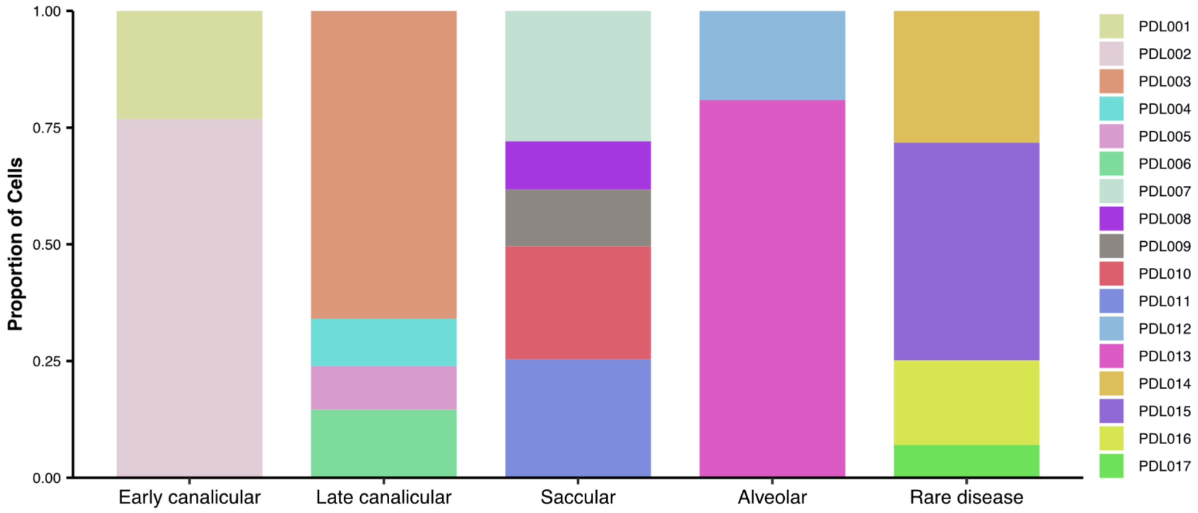


Supplemental Fig. S3:

A stacked bar chart showing the proportion of cells from each individual within each category.

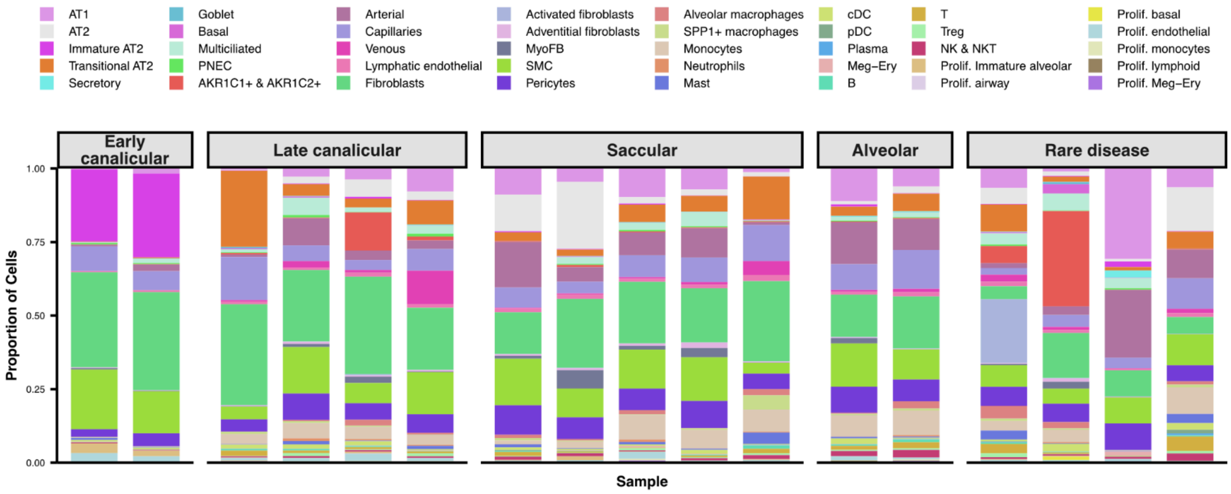


Supplemental Fig. S4:

A stacked barchart highlighting the proportion of each cell type across every sample in this cohort.

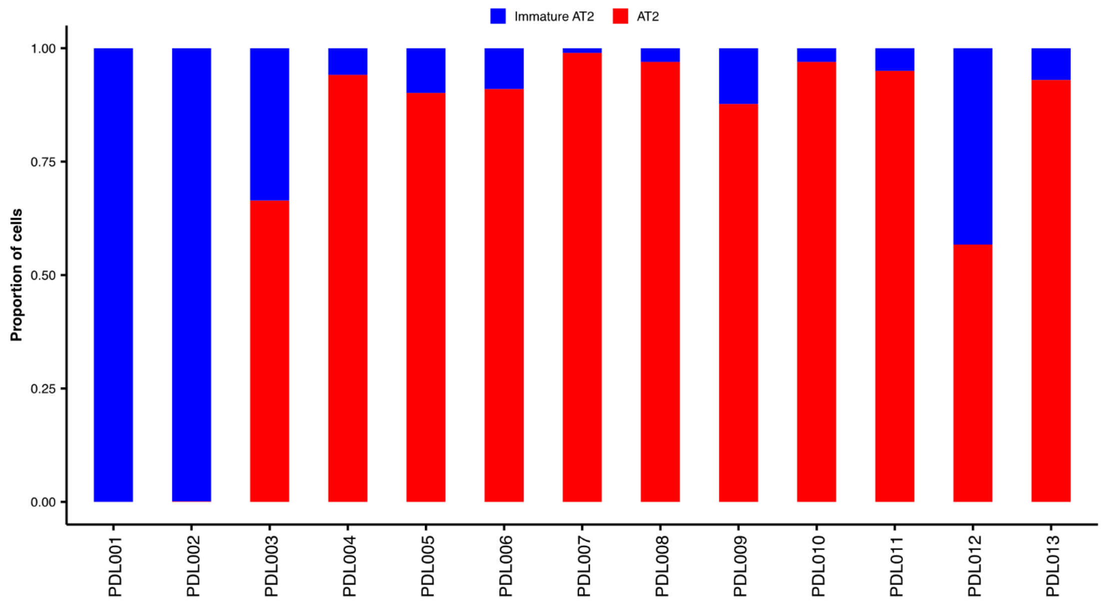


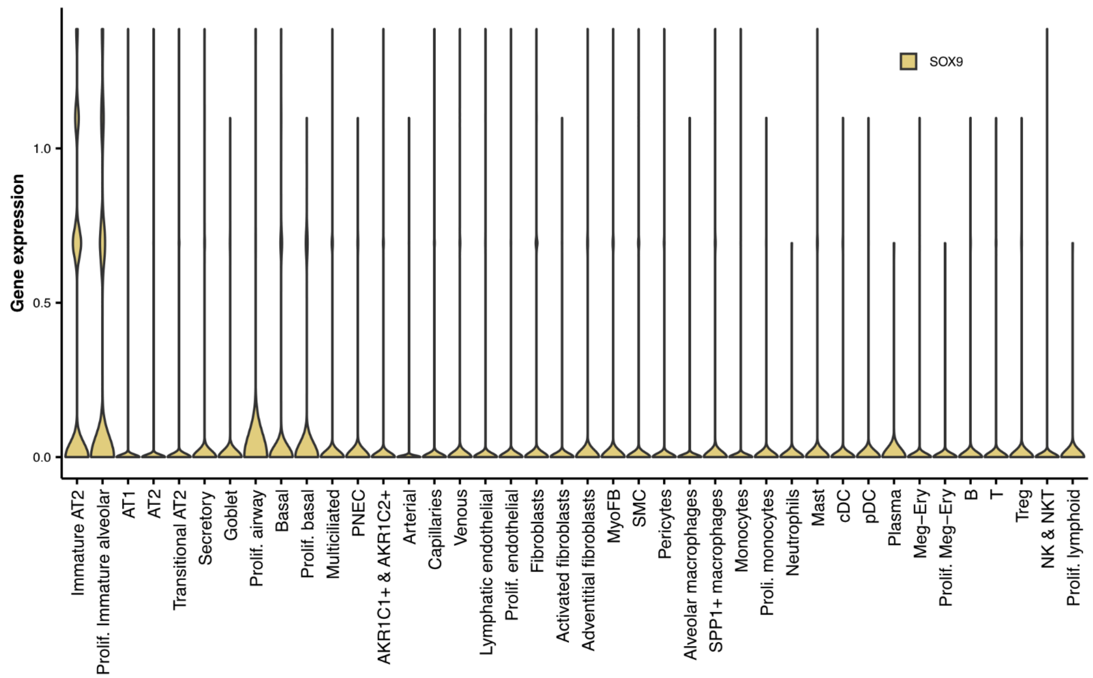


Supplemental Fig. S5:

a) A stacked bar chart showing the proportion of immature AT2 and AT2 cells across each individual, not including the rare diseased individuals. b) Violin plots showing the expression of *SOX9* across all cell types in cells from the same individuals used in (a).

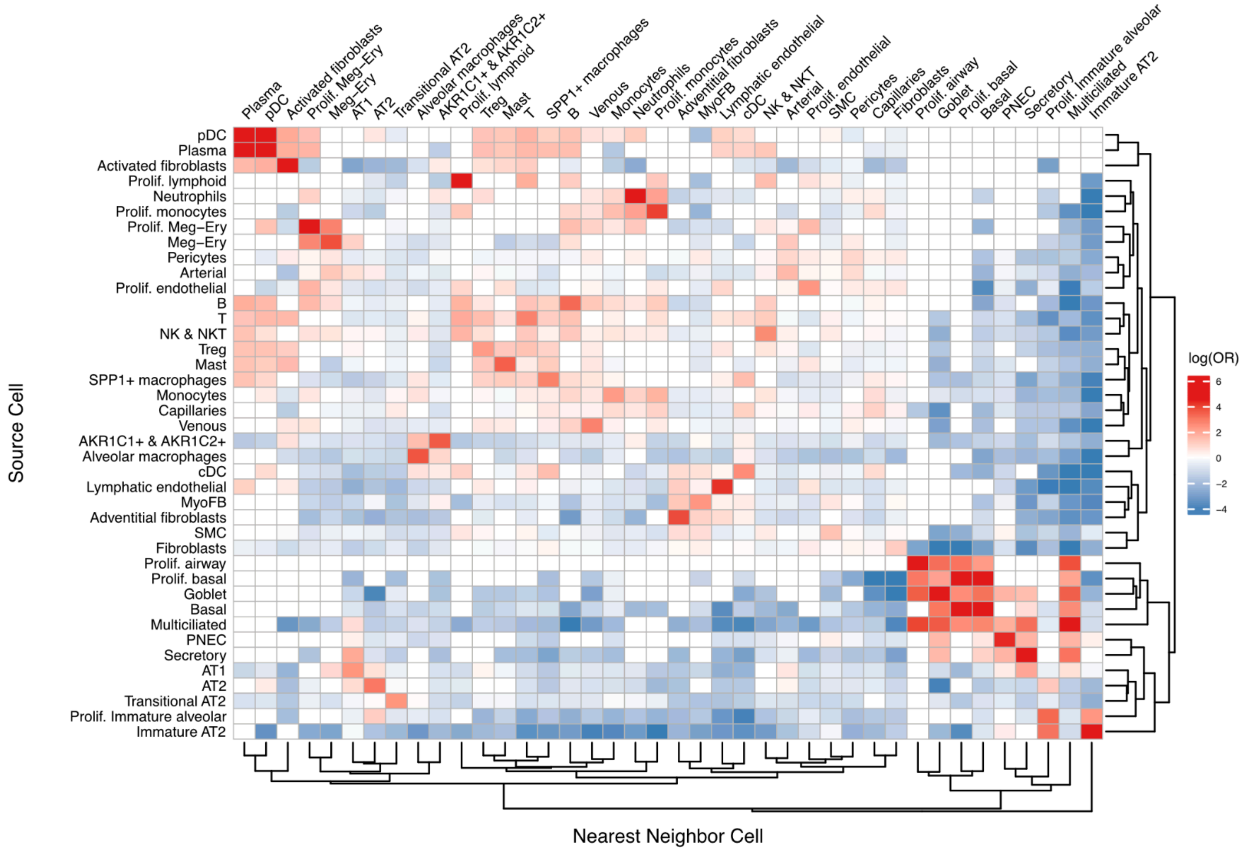


Supplemental Fig. S6:

Cell proximity results across all samples. This heatmap shows the log odds ratio (log(OR)) for each cell type pair, calculated with a Fisher's Exact test (**see Methods**). Positive values (red) indicate that the cells in that cell type pair are more likely to be located close to one another, while negative values (blue) indicate that those cells are less likely to be located close to each other. Non-significant results (FDR > 0.1) are coded as 0. See Supplemental figures 7-14 for cell proximity results across categories and specific samples representing rare diseases.

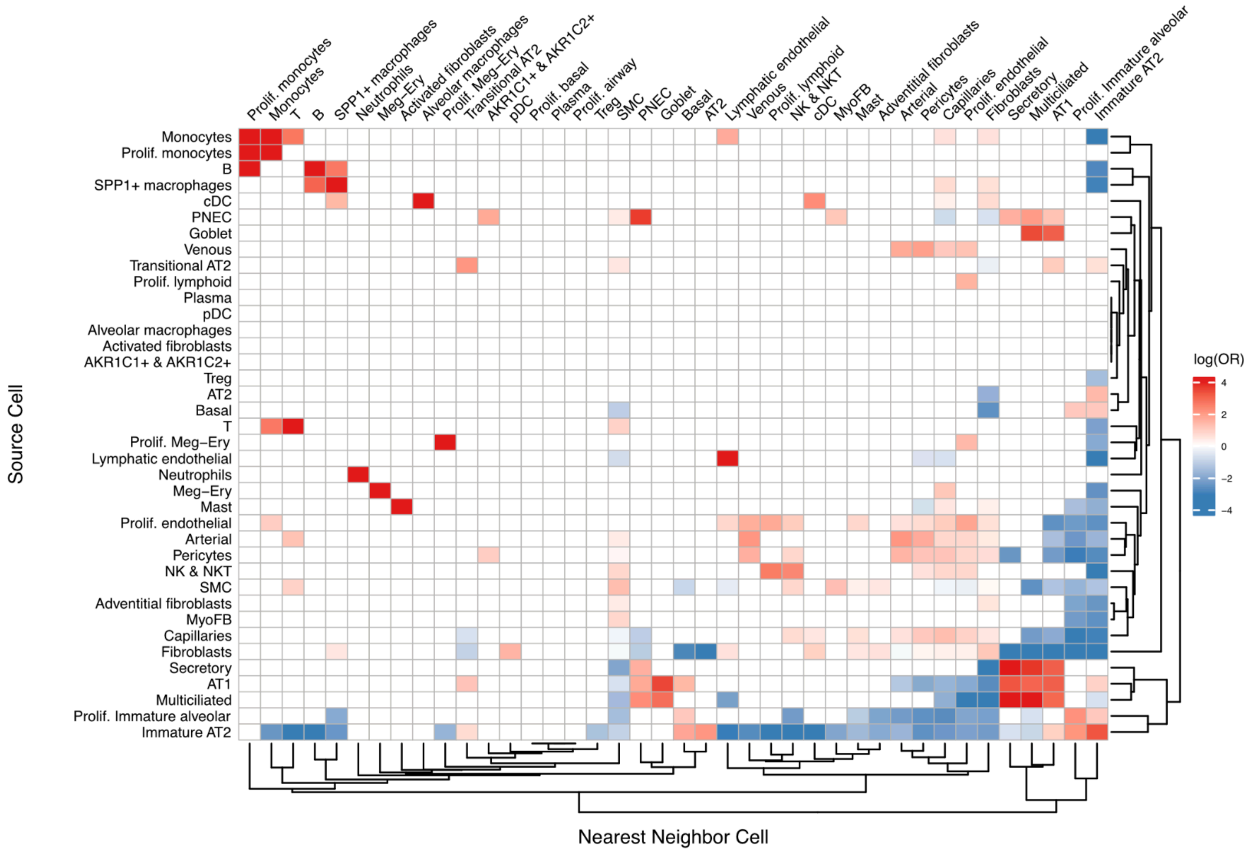


Supplemental Fig. S7:

Cell proximity results across all samples grouped into “early canalicular”.

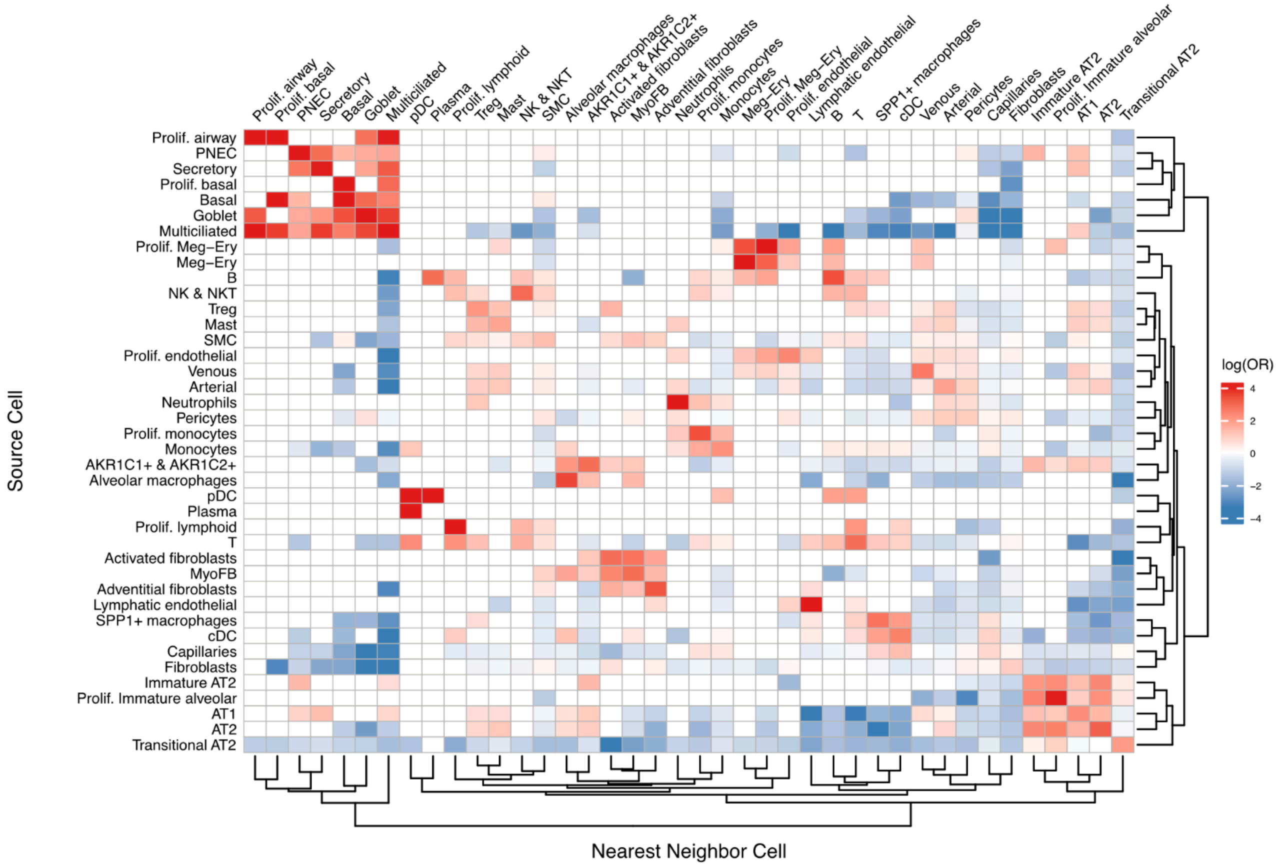


Supplemental Fig. S8:

Cell proximity results across all samples grouped into “late canalicular”.

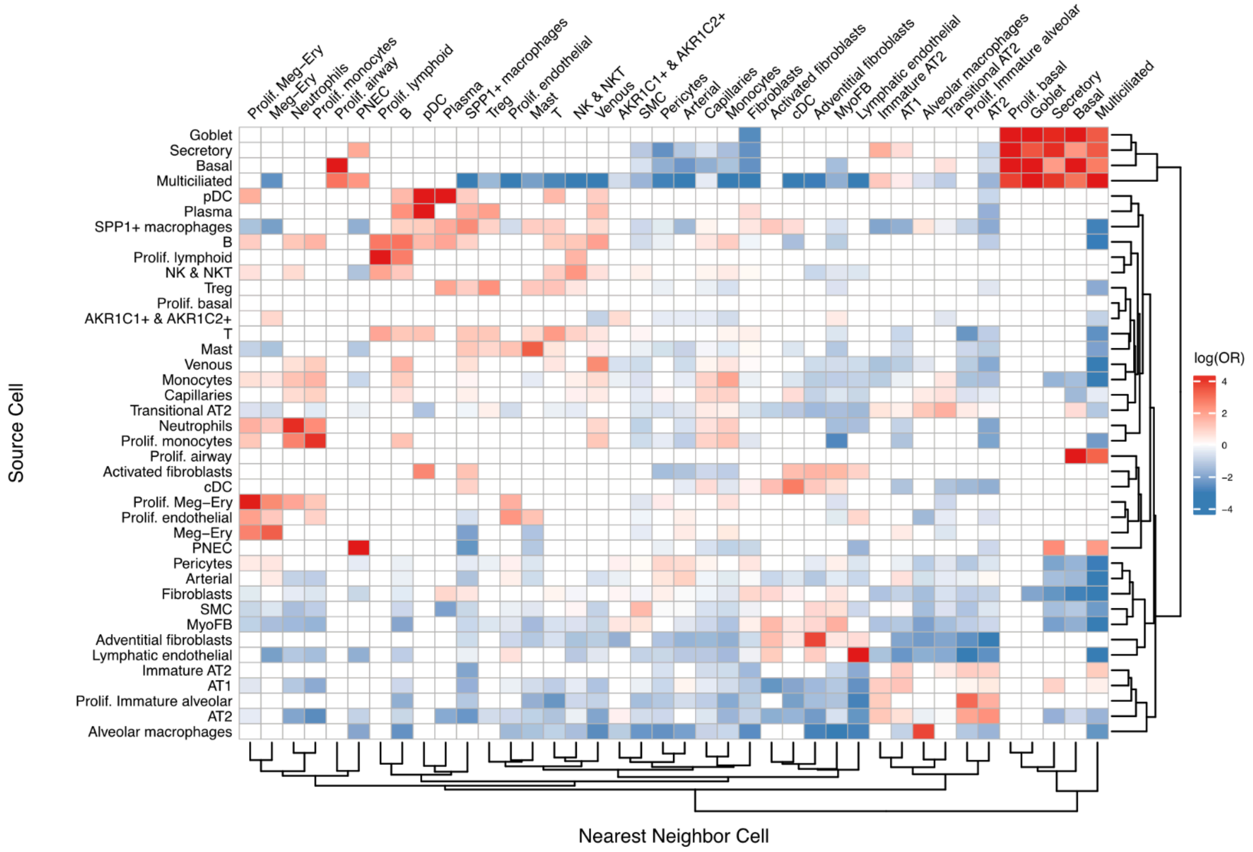


Supplemental Fig. S9:

Cell proximity results across all samples grouped into “saccular”.

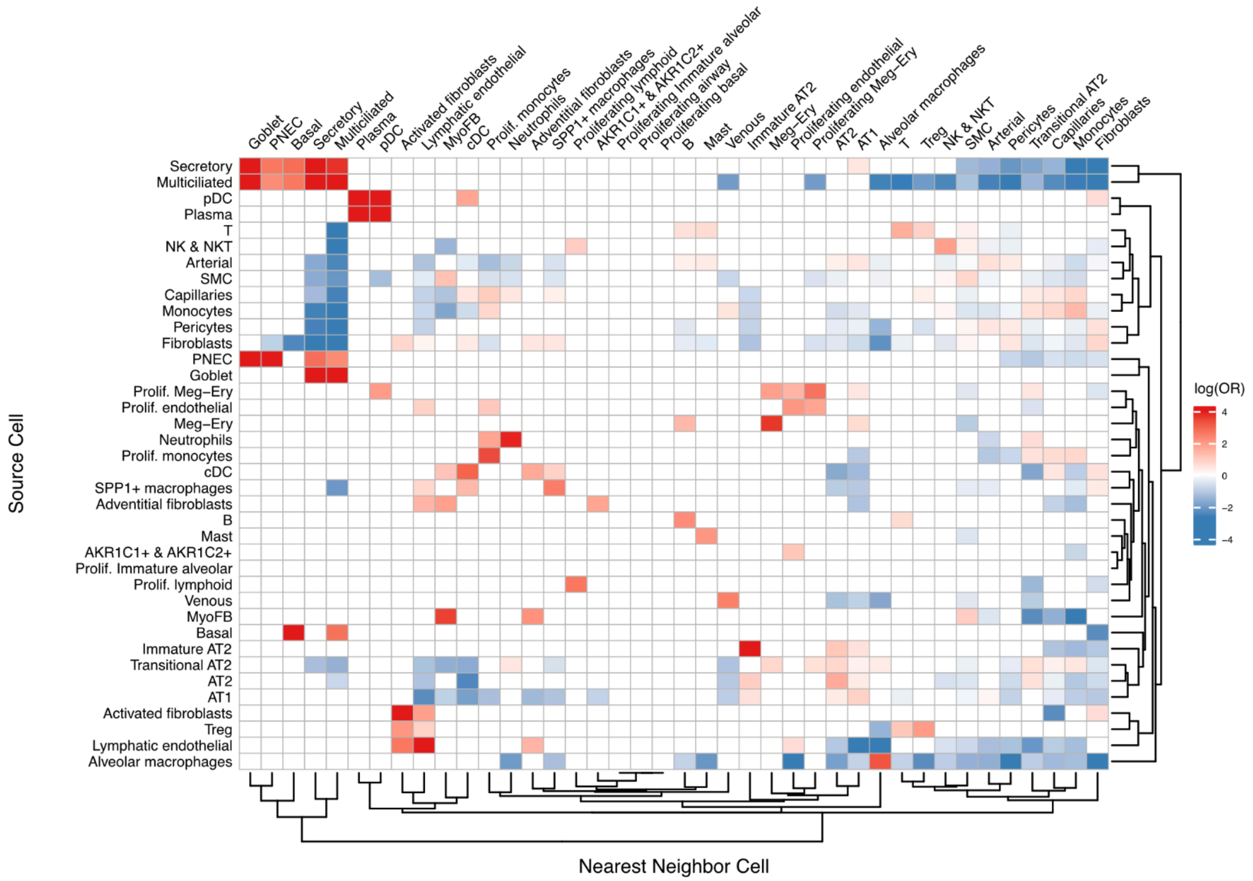


Supplemental Fig. S10:

Cell proximity results across all samples grouped into “alveolar”.

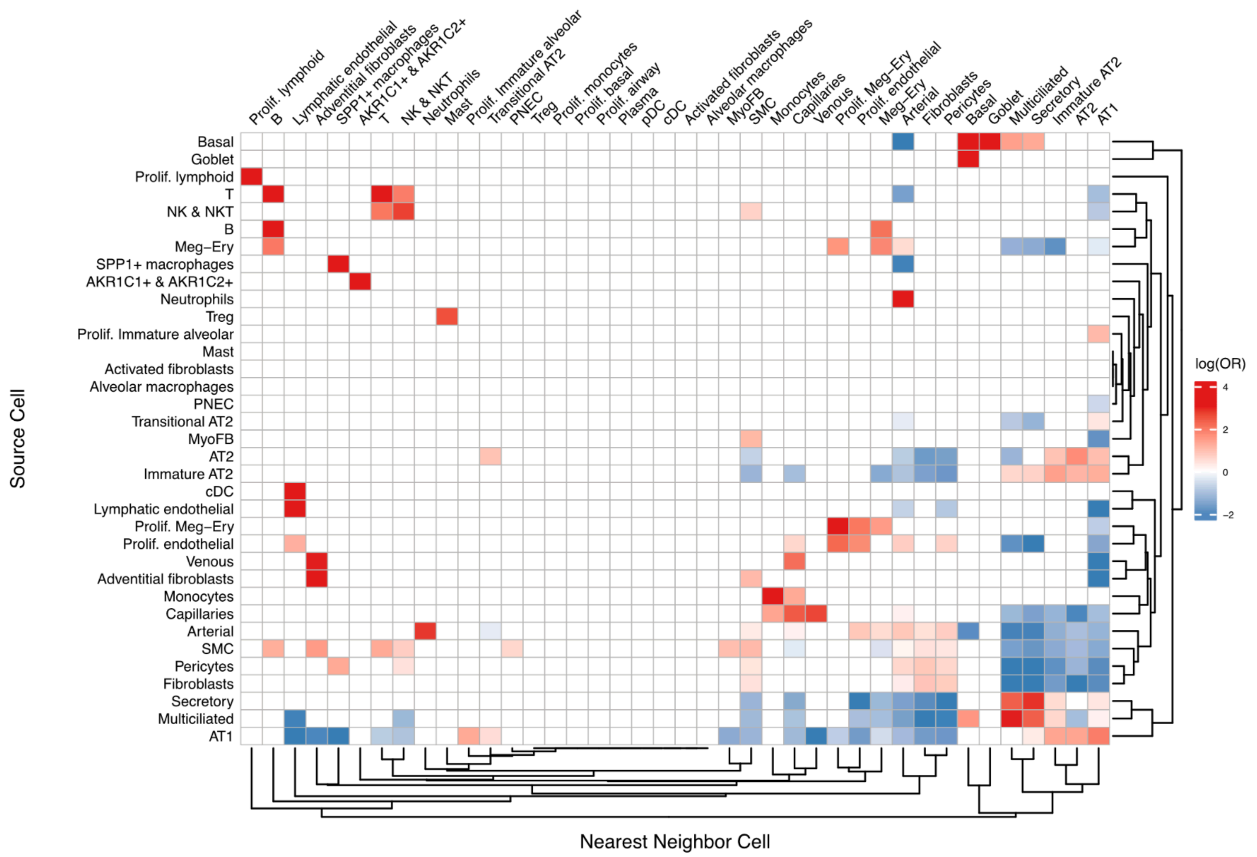


Supplemental Fig. S11:

Cell proximity results across one individual (PDL014) diagnosed with CHAOS syndrome who fell within the “rare disease” grouping.

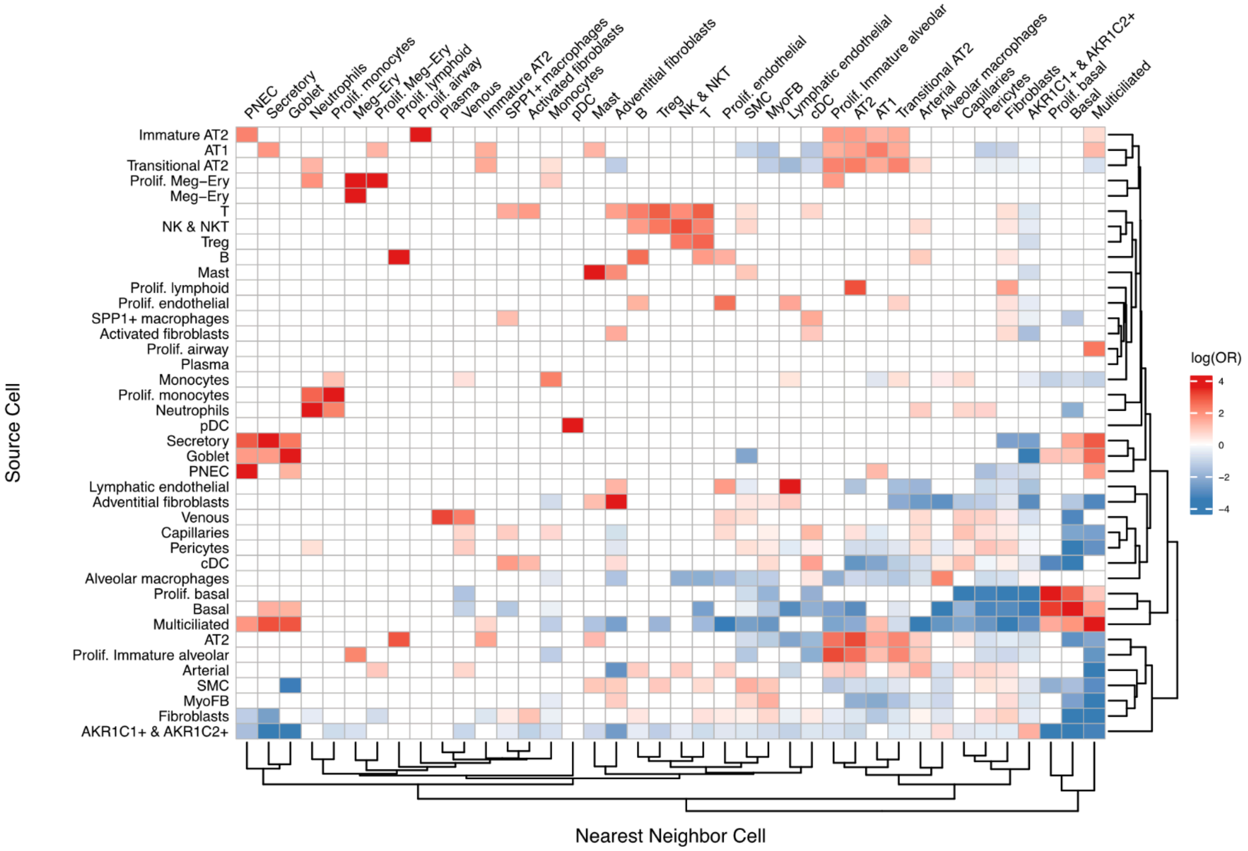


Supplemental Fig. S12:

Cell proximity results across one individual (PDL015) diagnosed with pulmonary hypoplasia and dysplastic kidneys who fell within the “rare disease” grouping.

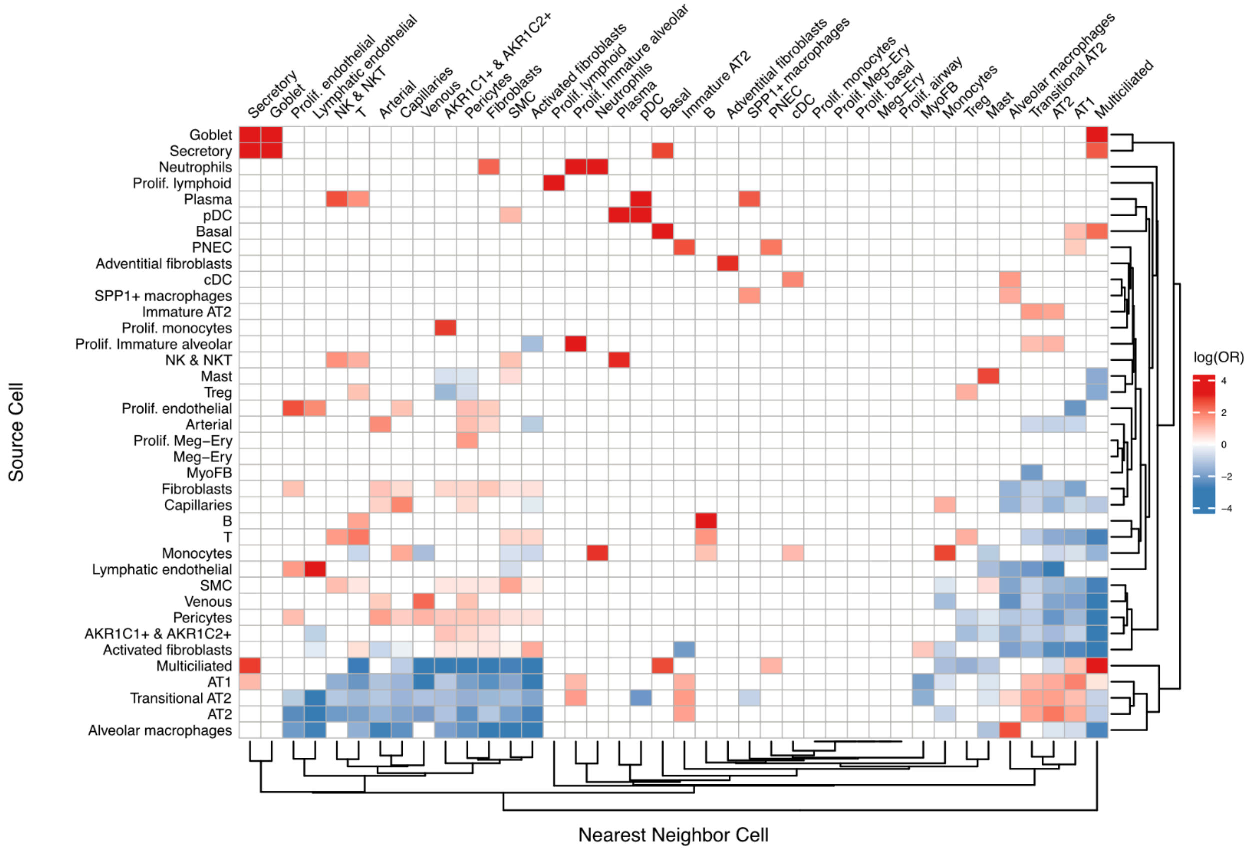


Supplemental Fig. S13:

Cell proximity results across one individual (PDL016) diagnosed with ARB fetopathy and severe BPD who fell within the “rare disease” grouping.

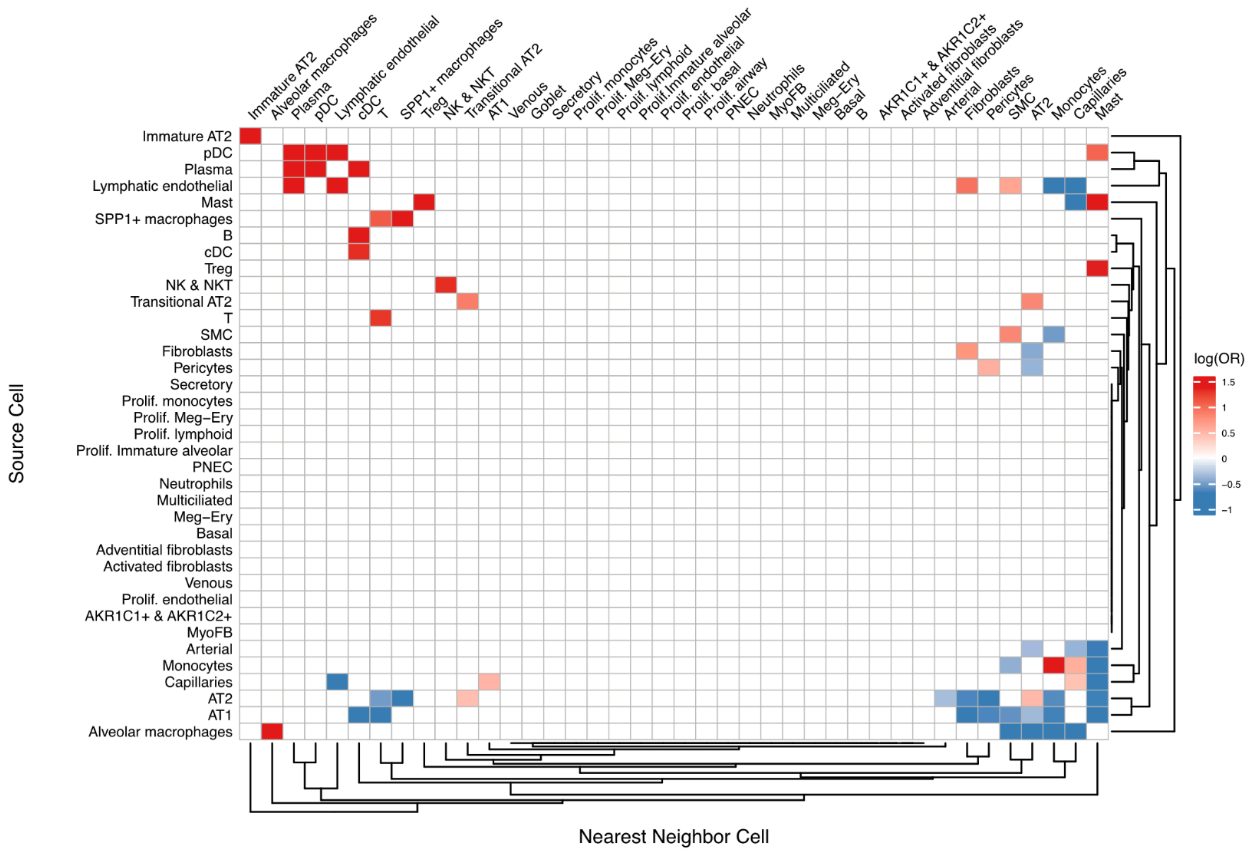


Supplemental Fig. S14:

Cell proximity results across one sample (PDL017) that was an explant of an adult aged 30 years living with BPD who fell within the “rare disease” grouping.

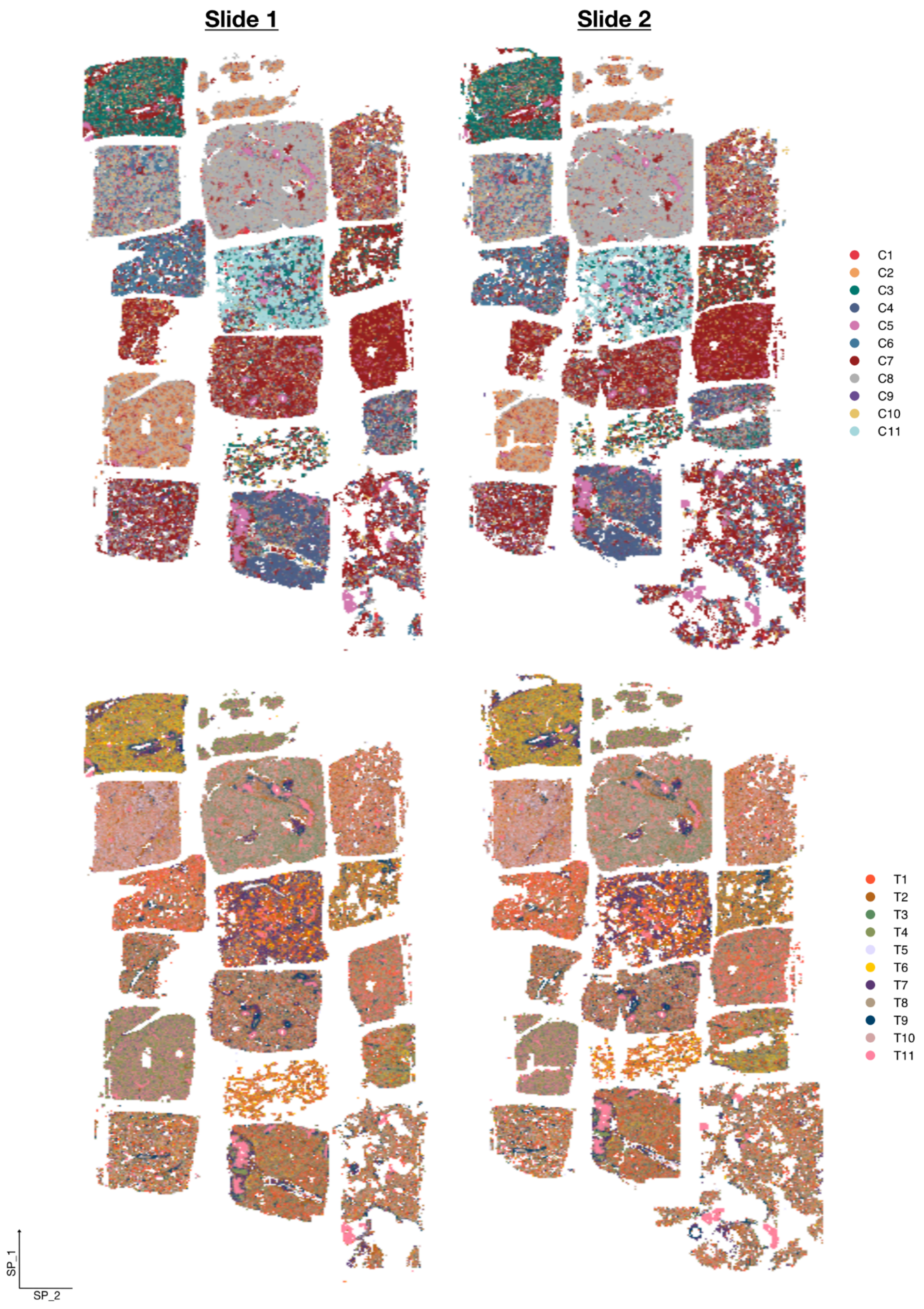


Supplemental Fig. S15:

Plots that show the structure of each tissue sample and showcase the distribution of niches across each sample, split across both slides. The top images are colored by the cell niches, and the bottom are colored by the transcript niches.

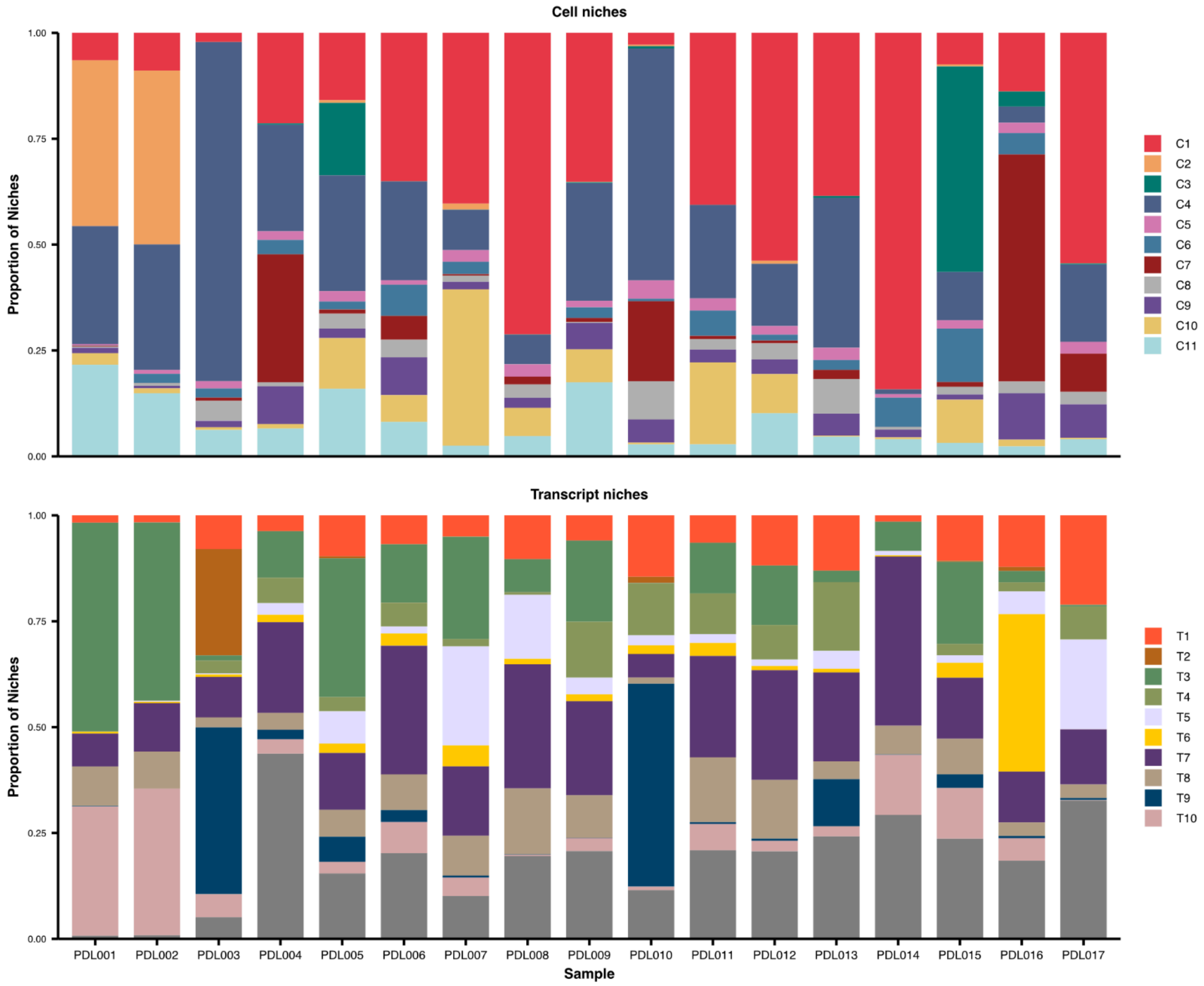


Supplemental Fig. S16:

Stacked bar plots show the composition of cell niches (top) and transcript niches (bottom) split across each individual.

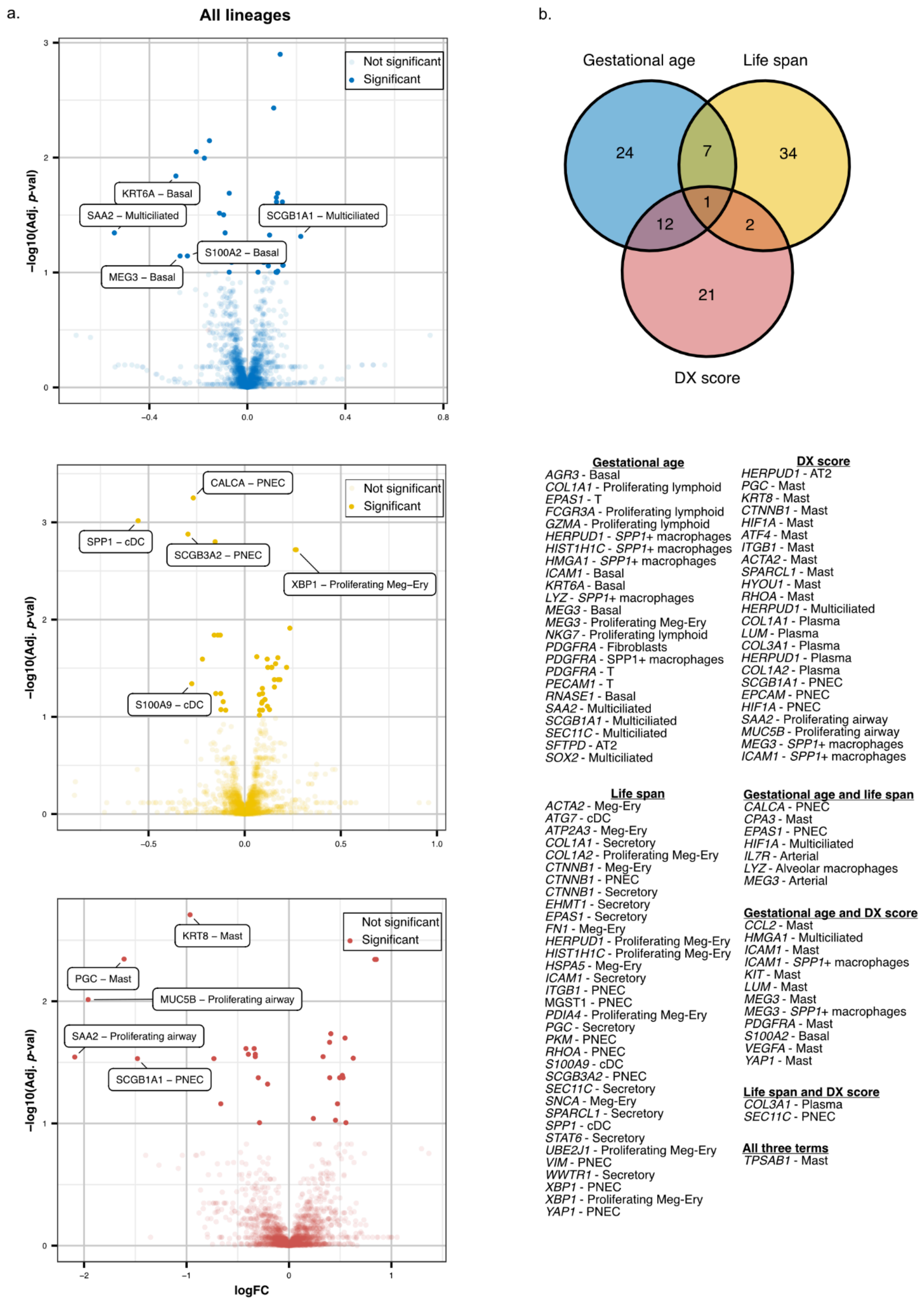


Supplemental Fig. S17:

a) Volcano plots showing the top 5 gene-cell type pairs in all lineages with the highest logFC per gestational age (blue), life span (yellow), and disease severity score (red). LogFC values in both were generated by a linear model we built that utilized the Limma package in R (**see Methods**). b) A venn diagram showing the overlapping gene-cell type pairs across all three terms within all lineages.

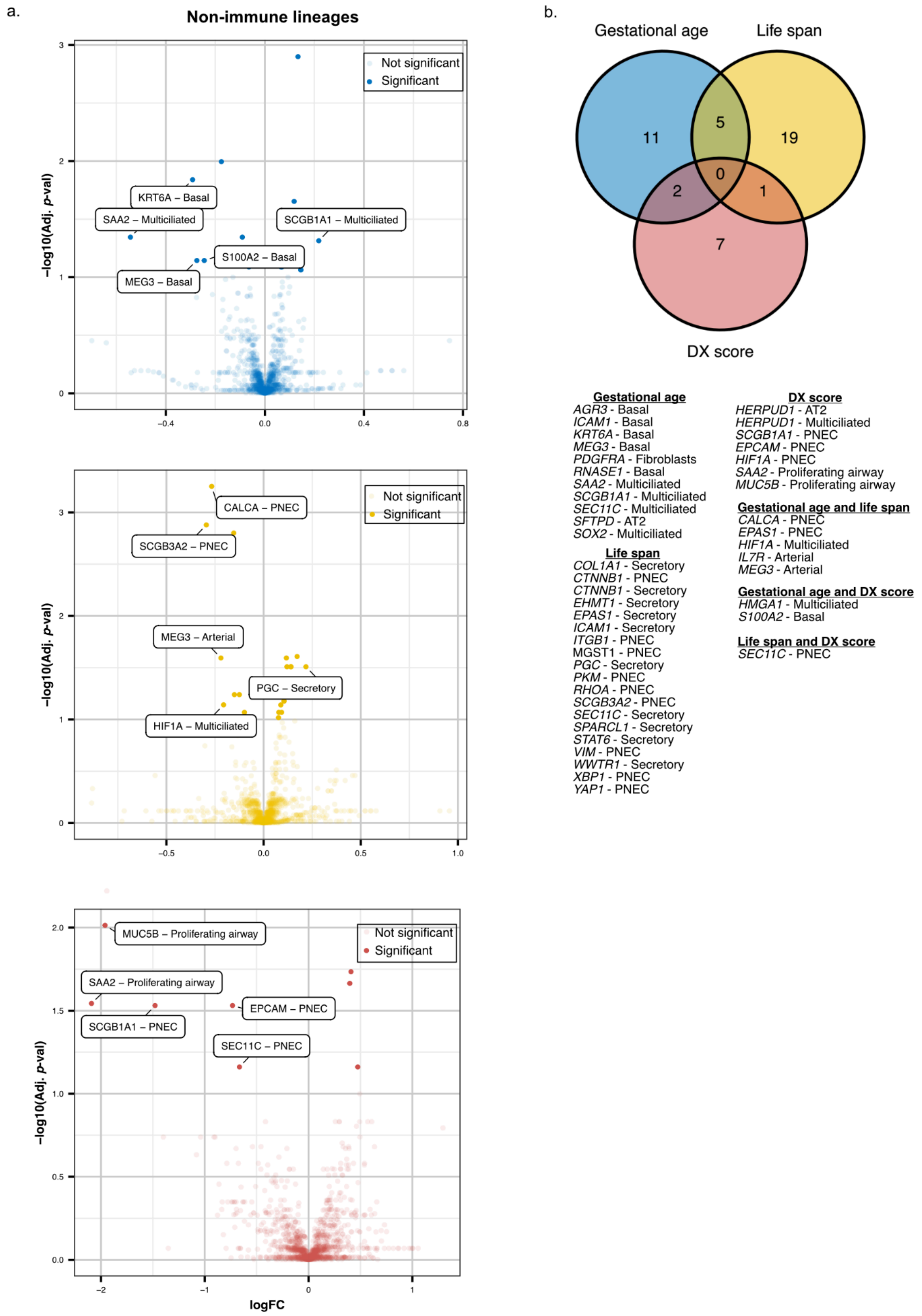


Supplemental Fig. S18:

a) Volcano plots showing the top 5 gene-cell type pairs in epithelial, endothelial, and mesenchymal lineages with the highest logFC per gestational age (blue), life span (yellow), and disease severity score (red). LogFC values in both were generated by a linear model we built that utilized the Limma package in R (**see Methods**). b) A venn diagram showing the overlapping gene-cell type pairs across all three terms within the epithelial, endothelial, and mesenchymal lineages.
